# Low-level features predict perceived similarity for naturalistic images

**DOI:** 10.1101/2024.08.15.607867

**Authors:** Emily J A-Izzeddin, Thomas SA Wallis, Jason B Mattingley, William J Harrison

## Abstract

The mechanisms by which humans perceptually organise individual regions of a visual scene to generate a coherent scene representation remain largely unknown. Our perception of statistical regularities has been relatively well-studied in simple stimuli, and explicit computational mechanisms that use low-level image features (e.g., luminance, contrast energy) to explain these perceptions have been described. Here, we investigate to what extent observers can effectively use such low-level information present in isolated naturalistic scene regions to facilitate associations between said regions. Across two experiments, participants were shown an isolated standard patch, then required to select which of two subsequently presented patches came from the same scene as the standard (2AFC). In Experiment 1, participants were consistently above chance when performing such association judgements. Additionally, participants’ responses were well-predicted by a generalised linear multilevel model (GLMM) employing predictors based on low-level feature similarity metrics (specifically, pixel-wise luminance and phase-invariant structure correlations). In Experiment 2, participants were presented with thresholded image regions, or regions reduced to only their edge content. Their performance was significantly poorer when they viewed unaltered image regions. Nonetheless, the model still correlated well with participants’ judgments. Our findings suggest that image region associations can be reduced to low-level feature correlations, providing evidence for the contribution of such basic features to judgements made on complex visual stimuli.

## 2. Introduction

Fundamental to our daily functioning is our ability to interpret the complexity of our visual surroundings, which in turn guides how we behave. When forming such interpretations, the majority of information is typically yielded from the current visual scene. Visual information is critical for efficient deduction of many task-relevant factors (Frazor & Geisler, 2006; Torralba & Oliva, 2003), such as determining if we have turned onto the right street on our way to work or finding the right product in a supermarket aisle. To perform such efficient visual interpretations, we inspect our environment through eye movements, allowing the high-definition fovea to fixate various areas of the scene (Akbas & Eckstein, 2017; Mirza et al., 2016; Noton & Stark, 1971). Based on such fixations, the visual system must construct a coherent representation of the scene we are viewing, integrating all relevant regions of visual space – a problem that has been identified throughout the literature for decades (Berens et al., 2021; Chen et al., 2023; Cohen et al., 1999; Goh et al., 2004; Hassabis & Maguire, 2007; Robertson et al., 2016; Singer & Gray, 1995; Steel et al., 2021; Treisman, 1998). However, it remains largely unclear how we perform this complex scene mapping process or, in the simplest case, how we associate two different regions of visual space with one another. Hence, in the current study, we investigated observers’ scene region association judgements on two isolated regions. Specifically, we were interested in how available information in isolated scene regions facilitates such associations, giving insight into whether and how such information may contribute to broader scene mapping processes.

Being able to perform scene region associations can have considerable implications for informing our current behaviour and goals. By scene region associations, we refer to observers’ ability to determine whether distinct subregions belong to the same scene. In performing scene region associations, we also inherently dissociate those regions from adjacent scenes/environments. Imagine standing in a central location of a house; you might be able to see, simultaneously, the living room, the dining room, the kitchen, and perhaps even other rooms through adjoining doors. This in and of itself can be considered to be one coherent scene (i.e., the inside of a house). However, if you were asked to find a jumper in the living room, it becomes functionally beneficial for you to focus on the smaller living room scene within the broader house scene. Therefore, to guide basic functions such as visual search, you need to delineate the boundary of the living room from other visible rooms and, critically, to associate the relevant regions of the living room with each other. Here, we emphasise the importance of *integrating* scene regions, as opposed to *segmenting* a scene. Such a process of refining the scope of the scene currently being considered demonstrates the flexible and task/goal-oriented process of scene-defining (Henderson & Hollingworth, 1999). However, the visual processing invoked to facilitate such scene-defining behaviour remains unclear.

There is an abundance of visual information available to us in naturalistic scenes, any subset of which we might conceivably use to associate regions of space (Henderson & Hollingworth, 1999). Heavy emphasis has been placed on the contribution of “semantic” visual information to scene processing, with consistent evidence for such high-level information being influential in visual object search and recognition (Bar, 2004; Bar & Ullman, 1993; Biederman et al., 1982; Brockmole et al., 2006; Brockmole & Henderson, 2006a, 2006b; Davenport & Potter, 2004; Eckstein et al., 2006; Friedman, 1979; Henderson et al., 1999; Henderson, 2003; Hidalgo-Sotelo et al., 2005; Hollingworth & Henderson, 2000; Loftus & Mackworth, 1978; Neider & Zelinsky, 2006; Palmer, 1975). However, given that semantic understanding of a scene follows from visual information, it is possible that semantics are, in some cases, redundant in scene processing. Instead, it is possible that lower-level visual information contributes uniquely to scene processing. Hence, beyond semantic visual information, there remains scope to further our understanding of scene processing by investigating the contribution of basic visual features.

Low-level features are a clear candidate for information we might use to associate regions of space. Here, low-level features are conceptually defined as any image information that does not convey semantic meaning (Neri, 2014). Operationally, we define low-level features as any information that can be computed from simple oriented contrast filters. Low-level feature relationships between regions of space have been explored computationally (Field, 1987; Frazor & Geisler, 2006; Harrison, 2021; Simoncelli & Olshausen, 2001). Practically, such computations typically involve some form of image decomposition that measures particular image features, such as contrast energy, at different orientation/frequency bands. From this, image statistics for a specific image region can be calculated – for example, quantifying the prevalence of particular orientations. Resulting statistics can then be correlated with the statistics of a different image region to provide a measure of similarity between the two regions in question. Such similarity assessments have suggested that the strength of these relationships is largely dependent on the relative spatial position of the two regions being associated (**Fig. 1**; Field, 1987; Frazor & Geisler, 2006; Harrison, 2021; Simoncelli & Olshausen, 2001). For example, there is a clear influence of spatial separation, whereby there tend to be stronger relationships between spatially proximal regions of space than more distant regions. Similarly, due to the over-representation of cardinal orientations relative to obliques in nature, low-level feature relationships will be stronger with cardinal separation axes than oblique separation axes (Coppola et al., 1998; Essock et al., 2003; Girshick et al., 2011; Hansen et al., 2003; Hansen & Essock, 2004; Harrison, 2021; Keil & Cristóbal, 2000). Hence, the computational capacity to associate regions of space based on low-level feature correlations is well-established.

**Figure 1.**
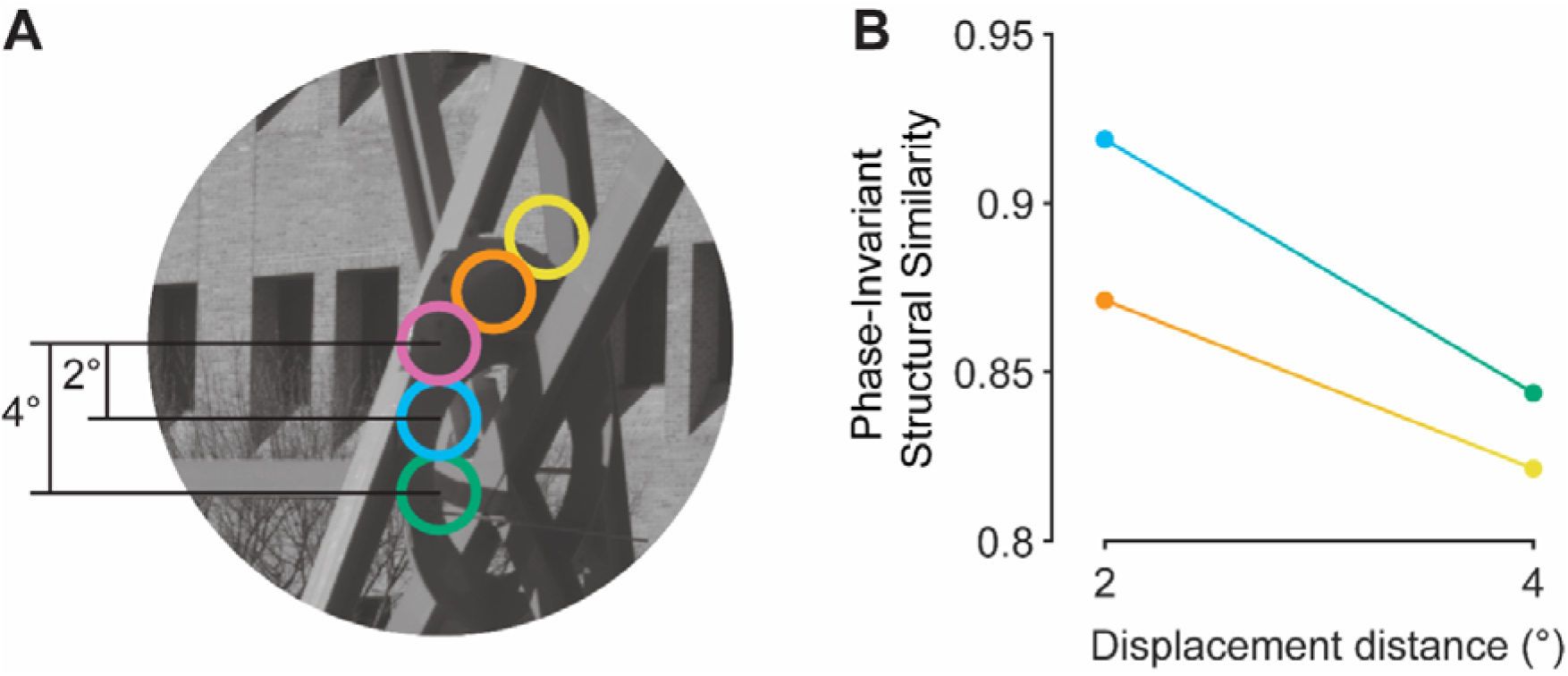
The impact of separation on image region correlations. **A)** The similarity between any two image regions can be measured using a variety of image-computable metrics. These image regions can relate to each other spatially according to different separation manipulations. For example, we can vary separation distance, comparing spatially proximal (e.g., orange and blue) vs distant (e.g., yellow and green) regions. Note, proximal vs distant are relative terms in this case. Alternatively, we can vary separation axis, comparing cardinally (i.e., blue and green) vs obliquely (i.e., orange and yellow) offset regions. These separations will have systematic impacts on the similarity between the pink and comparison region, as indicated on the right. **B)** One method for measuring similarity between image regions is to compute “phase-invariant structural similarity” (Sebastian et al., 2017), which quantifies the spatial similarity of the patches’ content, while ignoring the absolute luminance values of the pixels (see **Section 3.7.2., Generalised linear multilevel modelling**, for more detail). Here, the actual phase-invariant structural similarity between the pink region and other regions of interest is plotted on the y-axis, with separation distance on the x-axis, and separate lines for cardinal (blue and green) vs oblique (orange and yellow) separations. Spatially proximal regions (2°; orange and blue) have a higher similarity score than distant regions (4°; yellow and green). Additionally, higher similarity scores are observed for cardinal separations compared with obliques.

Beyond computational investigations, there has been extensive research into our sensitivity to basic visual information. Key insights into visual sensitivity have emerged from work investigating perceptual priors (expectations rooted in the statistical regularities we observe in nature) for such basic visual information and their subsequent impact on perception (Series & Seitz, 2013; Summerfield & Egner, 2009). For example, we have a greater sensitivity to cardinal orientations as compared with obliques (Appelle, 1972; Berkley et al., 1975; Campbell et al., 1966; Dakin, 2001; Dakin et al., 2009; Dakin & Watt, 1997; de Gardelle et al., 2010; Emsley, 1925; Girshick et al., 2011; Henderson & Hollingworth, 1999; Pratte et al., 2016; Westheimer & Beard, 1998) due to a prior that reflects our greater exposure to cardinal orientations in nature as compared with obliques (Coppola et al., 1998; Essock et al., 2003; Girshick et al., 2011; Hansen et al., 2003; Hansen & Essock, 2004; Harrison, 2021; Keil & Cristóbal, 2000). However, while observers are clearly able to make perceptual judgements based on such low-level information, they have typically been probed with these visual features presented in isolation (e.g., oriented gratings or bars; Carandini, 2005; David et al., 2004; Olshausen & Field, 2005; Sonkusare et al., 2019). Therefore, documented judgements on basic visual features are not necessarily representative of our judgements on the complex visual information conveyed by a naturalistic scene. Hence, the question remains as to whether we are sensitive to low-level information to the extent that it can facilitate naturalistic image region associations.

Previous investigations have sought to understand the association between low-level feature distributions and processing in more naturalistic stimuli. For example, changes in low-level image statistics, such as edge density, phase, and contrast, have been shown to predict target detection in photographic images (Bex et al., 2009; Bex, 2010; Rideaux et al., 2022; Sebastian et al., 2017, 2020; Spehar & Taylor, 2013; Wallis & Bex, 2012). Further, aesthetic preferences and complexity judgements on images seem to be aligned with stimuli that more closely reflect the typical underlying image statistics we experience in nature (Aks & Sprott, 1996; Knill et al., 1990; Spehar et al., 2003; Sprott, 1993; Taylor et al., 2011; Viengkham et al., 2022; Viengkham & Spehar, 2018). Such studies suggest that the low-level features present in naturalistic images are able to influence our perceptual experience.

In the present study, across two experiments, we investigated observers’ capacity to perform naturalistic scene region associations based on low-level visual features. Participants viewed small image patches, windowed from broader naturalistic images, and indicated which two of three were drawn from the same broader scene (**Fig. 2A**). By windowing the patches, we removed broader contextual information that could otherwise have been exploited to disambiguate the relationship between the patches. In this way, we encouraged participants to rely on the available low-level information (A-Izzeddin et al., 2022). Importantly, patches drawn from the same broader scene were selected based on various combinations of separation distances and azimuths, thereby altering their low-level feature relationships (**Fig. 2B**; Field, 1987; Frazor & Geisler, 2006; Simoncelli & Olshausen, 2001) and allowing us to investigate the impact of spatial separations on image region association judgements. Crucially, we used computational modelling to predict observers’ capacity to perform naturalistic scene region associations based on low-level visual feature correlations.

**Figure 2.**
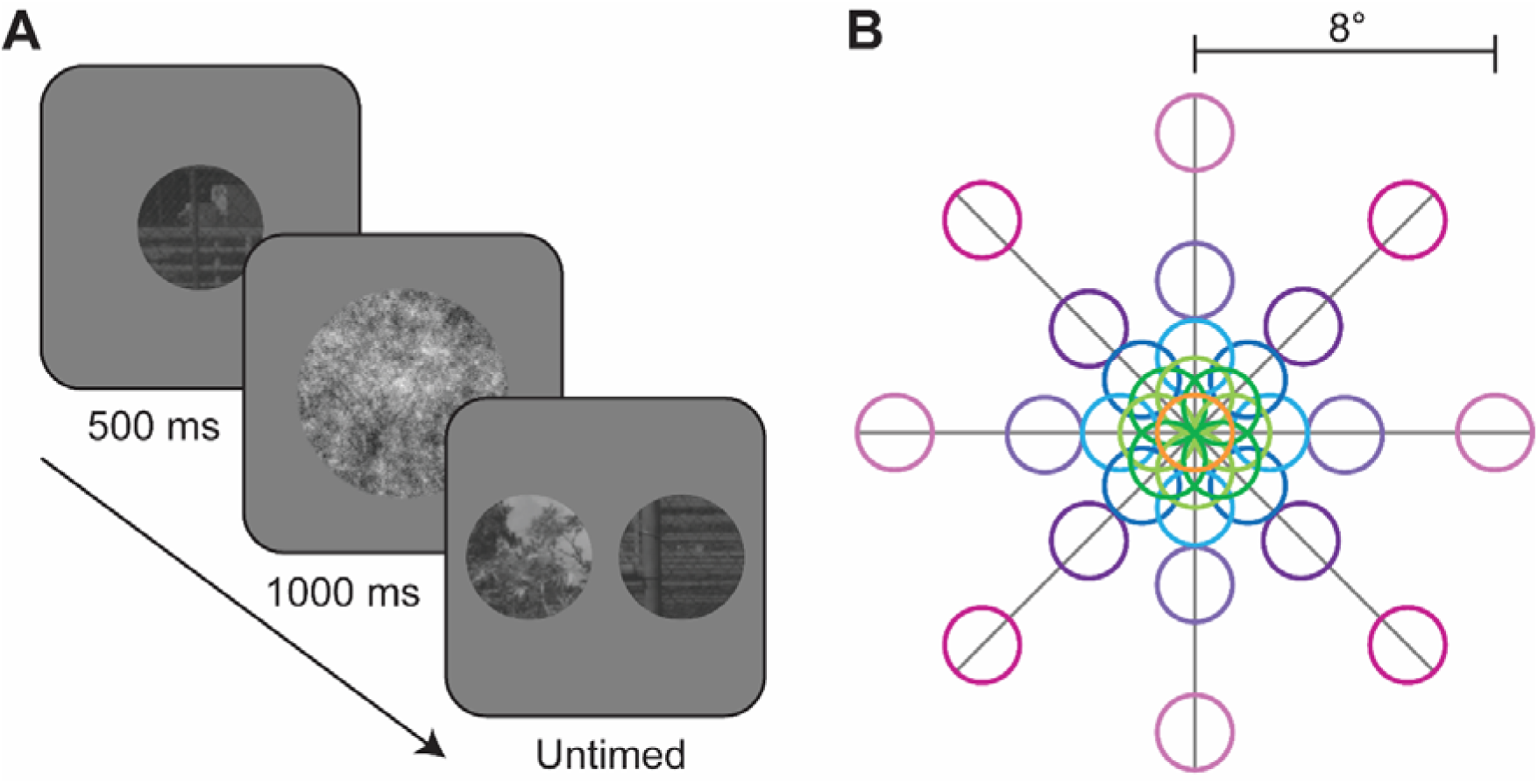
Experiment 1 paradigm overview. **A)** Schematic of general trial structure. On each trial, participants viewed a central fixation point (not pictured) for 700 ms, followed by the standard patch for 500 ms, a random pink noise mask (not to scale) for 1000 ms and, finally, the target (in this case, right) and foil (in this case, left) patches until a response was made. **B)** To-scale schematic of possible target/foil spatial locations relative to the standard patch (orange, also the location of the 0° separation condition), manipulating separation distance (0°, 1°, 2°, 4°, and 8°) and azimuth (0°, 45°, 90°, 135°, 180°, 225°, 270°, and 315°).

## 3. Methods

### 3.1. Participants

Two experiments were conducted, and each included data from 20 participants, aged 18-34. Participants were recruited through The University of Queensland’s Psychology Research Participation Scheme, facilitated by Sona. All participants were naïve to the purpose of the experiments. Ethics approval was granted by The University of Queensland (Medicine), Low & Negligible Risk Ethics Sub-Committee.

### 3.2. General task design

A schematic for a typical trial is shown in **Figure 2A**. Participants were first shown a fixation point in the centre of the screen. The fixation point was followed by a “standard” patch, which was an image region cropped from a larger photograph. Following the standard patch, participants saw a random pink noise mask, followed by two new image patches – a “target” and a “foil”. The target patch was another image region cropped from the same larger photograph as the standard, and the foil was cropped from an entirely different photograph. Participants were asked to indicate which of the two (target vs foil) came from the same larger photograph as the standard using the arrow keys on a standard computer keyboard. Participants did not receive feedback.

### 3.3. Stimuli

Stimuli were generated in the same manner across all experiments unless specified. Digital natural images were taken from a database of high-resolution 4256 x 2836-pixel colour photos, cropped to 104 evenly-spaced constituent 1080 x 1080- pixel regions (henceforth “source images”; Burge & Geisler, 2011). For both experiments, each trial comprised of one standard, one target, and one foil stimulus. For each trial, two unique source images were selected – one as the source for the standard and target patches, and the other as the source for the foil. The 104 source images cropped from a single original image in Burge and Geisler’s (2011) database contained overlapping content. Hence, the two source images on a given trial were selected to ensure they came from two different original images, preventing any overlap. Stimuli on each trial (i.e., the standard, target, and foil) were circular patches cropped from the selected 1080 x 1080 source images, subtending 2° of visual angle in diameter. All stimuli were converted to greyscale using MATLAB’s rgb2gray() function.

Standard patches were always cropped from the centre of the source image and were always presented to participants centrally. Original database images were taken of common scenes observed by the researchers at their university campus (Burge & Geisler, 2011). Hence, by cropping 1080 x 1080 regions of the original database photos and selecting one of these regions as the source image, any systematic ‘photographer’ bias which might bias the central region of the image (i.e., where the standard is drawn from) should have been eliminated. Target patches were cropped from one of 33 possible spatial locations relative to the standard patch (**Fig. 2B**). To further account for any potential photographer bias, foil patches were drawn from their source images at matched spatial locations to the target patches on each trial. The 33 possible spatial locations for the target and foil patches were specified combinations of distance (1°, 2°, 4°, or 8° of visual angle) and azimuth (0°, 45°, 90°, 135°, 180°, 225°, 270°, or 315°) separations relative to the standard patch (i.e., the centre of the image). Additionally, a 0° separation distance condition was included, where the target patch was identical the standard patch, to assess participants’ ability to perform the task when given identical region information. Target and foil patches were always presented simultaneously, separated either vertically or horizontally (counterbalanced across participants), with the side the target patch was on also balanced across trials.

### 3.4. Apparatus

For both experiments, stimuli were generated on a Dell Precision T1700 computer (running Windows 7 Enterprise) with the Psychophysics Toolbox (3.0.17; Brainard, 1997; Pelli, 1997) for MATLAB (R2020b). For the first 12 participants in Experiment 1 (including two excluded participants), stimuli were presented on a 32- inch Cambridge Research Systems Display++ LCD monitor with 1920 x 1080-pixel resolution, hardware gamma correction, and a refresh rate of 120 Hz. For the remainder of participants in Experiment 1 and all participants in Experiment 2, stimuli were presented on a 24-inch Asus VG428QE 3D monitor with 1920 x 1080-pixel resolution and a refresh rate of 120 Hz. This change was made because we moved labs during this project due to equipment availability constraints and had no consistent effect on participant performance (see **S.1**). A gamma correction of 2 was applied to the Asus display.

### 3.5. Experiment 1

Twenty-three participants completed Experiment 1. Upon visual inspection of individual data, three participants performed at chance in the 0° separation condition and were thus excluded from all other analyses. Each participant completed 1320 trials. All participants were shown the same set of stimuli, which comprised of 1320 unique standard/target source images and 1320 unique foil source images which were pseudorandomly selected from a bank of 9,361 potential images. Images were selected pseudorandomly to ensure unique source images were used for every stimulus, with no overlap in image content (per the source image generation technique used, described in **Section 3.3., Stimuli**), and no replacement after image selection. Trial order was randomised for each participant. Participants also completed 20 practice trials before starting the experiment, using randomly selected images not used in the experiment trials. On each trial, participants were shown a fixation point for 700 ms, followed by the standard patch for 500 ms, a random pink noise mask for 1000 ms and, finally, the target and foil patches (**Fig. 2A**). Target and foil patches were presented until participants indicated which one came from the same broader image as the standard. Importantly, participants were not told about the spatial separation manipulations used to generate the stimuli. Participants completed all trials in a single testing session split into 12 blocks, with self-paced breaks between each block.

### 3.6. Experiment 2

#### 3.6.1. Stimuli

To further investigate specific low-level feature contributions to scene region associations, Experiment 2 introduced an additional image manipulation. This new manipulation had three conditions, adapted from processing techniques used in Viengkham and Spehar (2022; **Fig. 3**). In the “full” patch condition, participants were shown unaltered foil/target patches (beyond the general processing outlined in **Section 3.3., Stimuli**). In the “threshold” patch condition, the median pixel value of the individual foil and test patches was calculated, and pixel values above this were set to white and pixel values below/equal to this were set to black. Finally, for the “edge” patch condition, source images were reduced to their edge content using MATLAB’s edge() function using the ‘Canny’ edge-finding method, with patches cropped from the resulting images. Regardless of the experimental condition, all standard images were left unaltered to avoid forcing participants to rely on certain types of cues when viewing the standard images. Rather, we sought to investigate whether participants had implicitly attended to such cues, benefiting their subsequent interpretation of processed test images.

**Figure 3.**
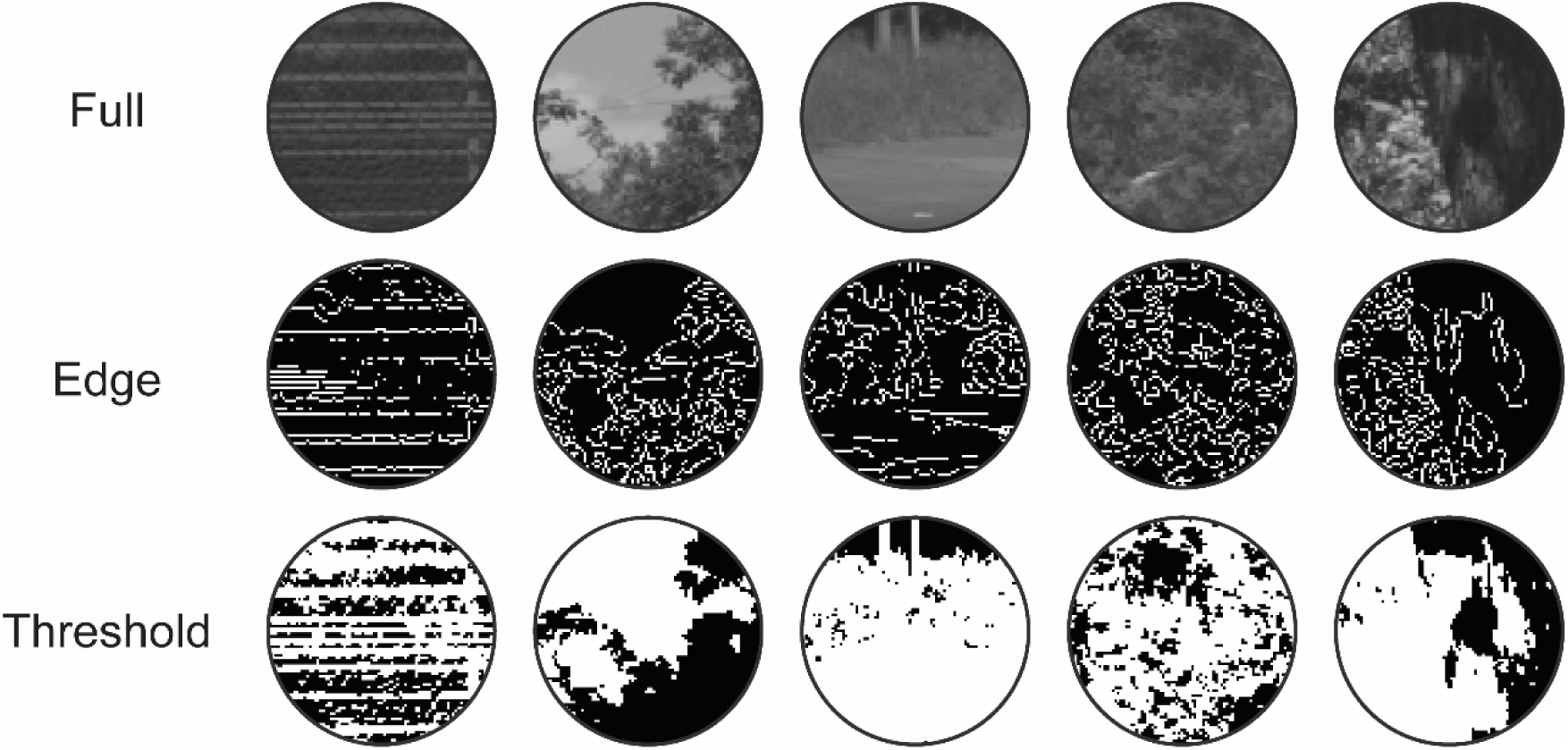
Example stimuli from each image processing condition in Experiment 2. Each row represents a different condition (i.e., full, edge, and threshold), with the same five source stimuli used for each row. Note: These conditions were only applied/compared for Experiment 2. Experiment 1 used full images only.

#### 3.6.2. Design

Participants (N=20; no exclusions) completed 1386 trials. All participants were shown the same set of stimuli, with the same stimuli also being used for each of the three image processing conditions. Hence, the stimulus set comprised of 462 unique standard/target source images and 462 unique foil source images, which were pseudorandomly selected from a bank of 9,361 potential images. Again, images were selected pseudorandomly to ensure unique source images were used for every stimulus, with no overlap in image content (per the source image generation technique used, described in **Section 3.3., Stimuli**), and no replacement after image selection. Trial order was randomised for each participant. Participants completed 20 practice trials before starting the experiment, using randomly selected images not used in actual experiment trials. On each trial, participants were shown a fixation point for 700 ms, followed by the standard patch for 500 ms, a random pink noise mask for 200 ms and, finally, were shown the target and foil patches (**Fig. 2A**). Target and foil patches were presented until a response was made with a button press, indicating which patch participants believed came from the same broader image as the standard. Again, participants were not made aware of the spatial separation manipulations used to generate the stimuli. Participants completed all trials in a single testing session split into 14 blocks, with self-paced breaks between each block.

### 3.7. Analyses

#### 3.7.1. General

Inferential statistics, where performed, were Bayesian analyses conducted in JASP, using response accuracy as the dependent variable.

#### 3.7.2. Generalised linear multilevel modelling

We modelled participants’ perceptual decisions under a generalised linear multilevel modelling (GLMM) framework, fit to all participants’ data in each experiment. The GLMM was implemented using MATLAB’s fitglme() function. Participants’ responses were coded as 0 or 1, to indicate incorrect and correct responses, respectively. The model was provided with predictors based on the same stimuli participants saw across both conditions – comprising of only full patches for Experiment 1, or combinations of full/edge/threshold patches for Experiment 2. Predictors were based on stimuli provided as square images, with the circularly windowed stimulus patch flanked by uniform mid-grey. Like participants, the model was not given access to information about the image-separation (in other words, separation distance and azimuth were not entered as predictors). Instead, predictors involving two image statistics based on low-level (i.e., image computable) information were included in the model: pixel-wise luminance root mean square (RMS) error and phase-invariant structural similarity, each of which is described below. We predicted the probability of a correct response as:

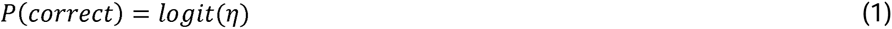

where *logit* represents the logit link-function, and:

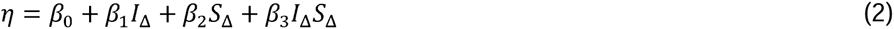

where *β*_0_ is the intercept term, *β*_1_ is the weight of the pixel-wise luminance difference, *I*_Δ_, *β*_2_ is the weight of phase-invariant structural similarity, *S*_Δ_, and *β*_3_ is the weight of the interaction *I*_Δ_*S*_Δ_. *I*_Δ_ and *S*_Δ_ are defined below. We implemented this model as a GLMM to partially pool coefficient estimates across observers and images, both of which were random effects in the model. The random effect of images refers to the unique image combination on a given trial. We describe this approach in more detail in Rideaux et al. (2022).

Our first predictor – pixel-wise luminance error – involved computing the RMS error between the standard patch and the target patch, as well as between the standard patch and the foil patch. We then determined which patch was most similar to the standard by calculating the difference score between these error scores, subtracting the foil luminance error from the target luminance error, giving *I*_Δ_.

Positive values of *I*_Δ_ indicate that the foil patch was more similar to the standard (in terms of pixel distances), whereas negative values of *I*_Δ_ indicate that the target patch was more similar to the standard. We finally log-scaled absolute *I*_Δ_ values, multiplied them by their original sign, and standardised by dividing each value of *I*_Δ_ by the standard deviation of all values of *I*_Δ_. We note here that *I*_Δ_ values were never 0, given these represent differences in the target and foil patches’ similarity scores. Hence, *I*_Δ_ could only be 0 if both the foil and target had the same similarity scores – this is only likely to occur should the foil and target be identical, which was never the case in the current study.

Our second predictor was phase-invariant structural similarity. A phase-invariant measure of structural similarity was included due to the likelihood of substantial luminance changes across a photo (e.g., some areas might be in direct sunlight and others in shade). Such variations in luminance could lead to very low pixel-wise luminance correlations, despite the underlying structure remaining the same. Hence, the phase-invariant structural similarity predictor quantified the spatial similarity of the patches’ spatial frequency and orientation content, while ignoring the absolute luminance values of the pixels (Sebastian et al., 2017). Per Sebastian et al. (2017), we first found the amplitude spectrum of the standard (A*_S_*), target (A*_T_*), and foil (A*_F_*) patches: for each patch, amplitude spectra were calculated by subtracting the mean of the patch, applying a fast Fourier transform (FFT), and taking the absolute complex value. To quantify the structural similarity between the standard and the target, as well as between the standard and the foil, we took the cosine of the vector angle between the amplitude spectra of the relevant test patch:

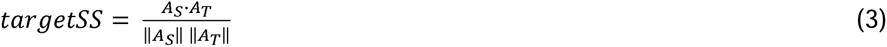

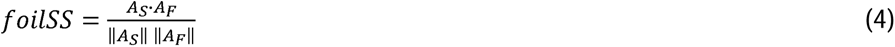

We then determined which patch was most similar to the standard by calculating the difference score between these structural similarity scores, subtracting foilSS from targetSS, giving *S*_Δ_. Positive values of *S*_Δ_ indicate that the target patch was more similar to the standard (in terms of structural similarity), whereas negative values of *S*_Δ_ indicate that the foil patch was more similar to the standard. We finally log-scaled absolute *S*_Δ_ values, multiplied them by their original sign, and standardised by dividing each value of *S*_Δ_ by the standard deviation of all values of *S*_Δ_.

We note that in Experiment 2, one trial in the edge-only condition resulted in target and foil images that were both devoid of any edges (i.e., two black patches were presented). As such, both patches yielded the same similarity scores for both predictors, leading to a difference score of 0, which resulted in an infinite value after undergoing log-scaling. We therefore removed trials with this combination of patches (across all image filtering conditions) from analyses – this was a total of three trials per participant, as each participant saw the same set of images, and each image filtering condition repeated the same source set of images.

To visualise the model fits (e.g., **Fig. 4 & 6-7, solid lines**), we generated predicted response probabilities for each trial. Predictions were yielded from the GLMM fit output, ‘F’, using ‘F.predict’. Note, in generating trial-by-trial predictions, we were able to partial predictions into separation distance/azimuth conditions with associated standard error of the mean estimates. To reiterate, however, separations were not used as model predictors.

**Figure 4.**
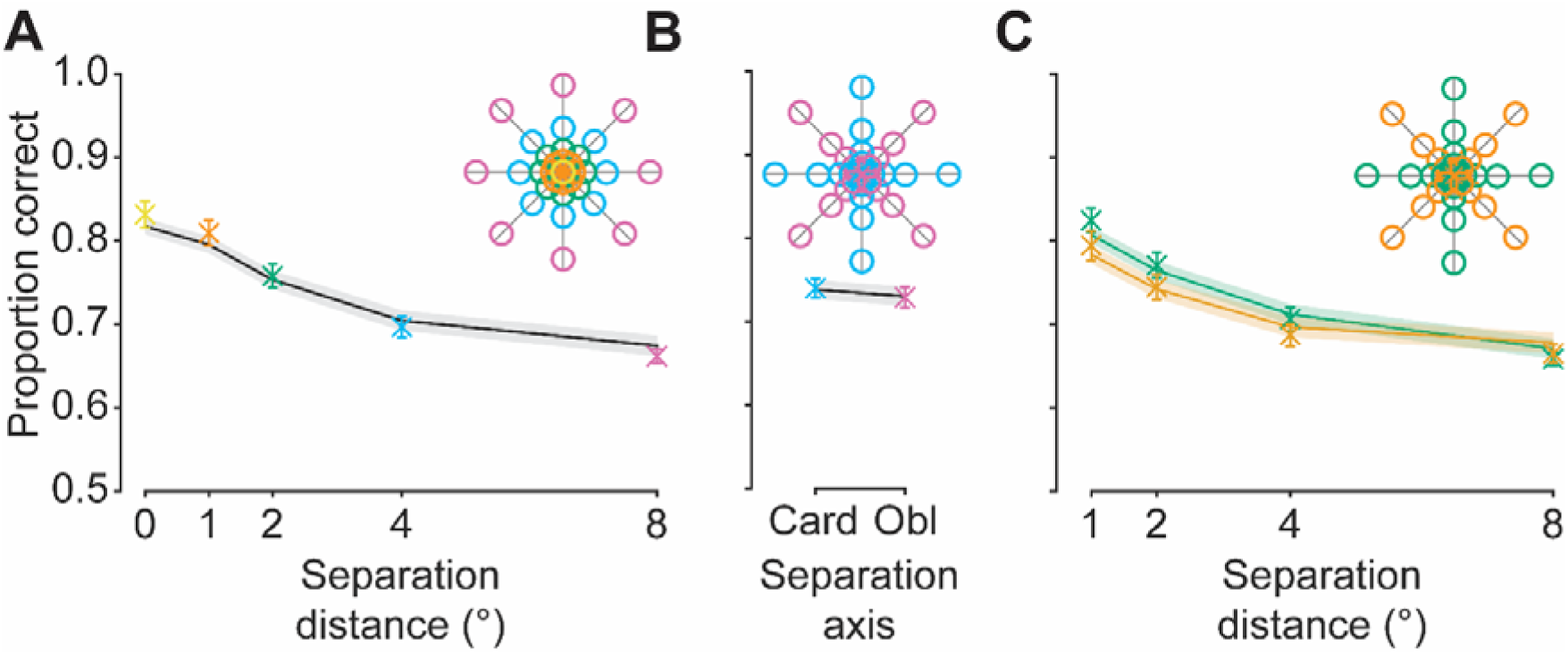
Experiment 1 behavioural and GLMM results. **A)** Effect of separation distance (x-axis) on the proportion of correct responses (y-axis). Data points are colour coded and are averaged across spatial locations of the same colour in the legend (inset). Solid line represents GLMM predictions based on luminance and structural similarity heuristics between the target and standard patch vs the foil and standard patch. **B)** Effect of separation axis (cardinal vs oblique) on the proportion of correct responses. **C)** The interaction between separation distance (x-axis) and separation axis (separate lines; see legend) on the proportion of correct responses. Error bars: ±1 SEM for participant responses (in some cases, standard errors are smaller than the point size). Shaded regions: ±1 SEM for trial-by-trial model predictions.

We implemented an alternative GLMM that was identical to the current model described above, however using log ratio scores instead of difference scores. Specifically, we calculated both pixel-wise luminance error and phase-invariant structural similarity scores as in the previous model comparing the target and foil to the standard. For each predictor, we then divided the target’s similarity score by the foil’s and took the log of this value. To avoid infinite values, we added a constant value of 10^-6^ to similarity scores before taking the log ratio. This was particularly relevant in the 0° separation condition, where pixel-wise luminance error for the target is 0. We found highly similar fits for this model as compared with the difference score model across both experiments, and therefore only report difference score model results.

We compared the performance of the full model (as described above) to alternative difference score models containing only the phase-invariant structural similarity main effect, or both the phase-invariant structural similarity and pixel-wise luminance difference main effects. Nested models were formally compared, with full output and comparison results presented in the supplemental materials for both Experiment 1 (**S.4**) and Experiment 2 (**S.14**). While we note that the full model was not the best-performing model in all comparisons, in the main body of the current text, we report the full model results for consistency.

## 4. Results

### 4.1. Experiment 1: Can similarity judgements be predicted from low-level features?

In Experiment 1, participants were presented with two image patches (a target and a foil) and selected which they judged to have been drawn from the same broader image/scene as a preceding standard patch (**Fig. 2A**). The spatial location that the target patch was drawn from relative to the standard was manipulated via distance (0°, 1°, 2°, 4°, or 8° of visual angle) and azimuth (0°, 45°, 90°, 135°, 180°, 225°, 270°, or 315°; **Fig. 2B**), but participants did not have knowledge of this spatial information. To assess performance, the proportion of correct responses (i.e., correctly identifying the target patch) was calculated for each condition. Based on known pixel-wise correlations between image regions and how they change with separation, we expected participants’ performance to decline with increasing separation distance, and that participants would perform better for cardinal separation axes than for obliques (Field, 1987; Frazor & Geisler, 2006; Simoncelli & Olshausen, 2001). We first describe the qualitative patterns of the data, and then describe the quantitative GLMM fits.

Behavioural data, as shown in **Figure 4**, were inspected by a 2 (separation axis: cardinal, oblique) x 4 (separation distance: 1°, 2°, 4°, 8°; 0° was not included as this condition could not interact with separation axis) Bayesian repeated measures ANOVA. As expected, participants’ performance decreased as the separation distance increased between the standard and the target, supported by extreme evidence in favour of an overall separation distance main effect (BF_10_ = 9.662x10^38^) and with post-hoc comparisons in favour of a difference in task performance between each individual separation distance condition (minimum BF_10_ = 893.056). Counter to expectations, however, there was no clear effect of separation axis on performance (BF_10_ = 0.528), with comparable accuracy levels for cardinal and oblique offsets (see **S.1** for individual data). Visually, there appears to be an interaction effect between separation axis and distance. Specifically, there appeared to be a benefit for cardinal offsets at the smallest distances that diminished in magnitude with increasing separation distance. This result pattern would suggest that information carried along the cardinal axes is only noticeably useful (beyond that observed for oblique axes) up to a certain separation distance, after which information conveyed by either separation axis is equivocal. However, there was equivocal evidence in favour of the alternative and null hypotheses in relation to the interaction (BF_10_ = 1.083). As such, there was no clear effect of separation axis on task performance.

When provided with two identical target patches (i.e., the 0° separation condition), participants’ response accuracy was worse than what would be expected if their performance in this condition depended only on a simple lapse rate. It is possible that this result is due to a proportion of memory-related errors and response errors, both of which should be distributed uniformly across separation conditions. Importantly, all participants show a very similar pattern of result across all separation distances, but with substantial differences in absolute performance across observers (see **S.1** for individual data). We further note that there was greater between-subject variance for the 0° separation condition as compared with the other separation distances (see **S.2**). As such, we tentatively suggest that the interaction between individual participant performance, as well as the increased variance observed for the 0°, were the primary contributors to the unexpectedly low performance observed for the 0° separation condition.

A GLMM was fit to the data to quantify performance using low-level, or image computable, information. Predictors were similarity metrics based on pixel-wise luminance and phase-invariant structure correlations. Raw pixel-wise luminance and phase-invariant structural similarity scores and their interactions with separation conditions were inspected independently of their use as predictors in the GLMM. Specifically, as separation distance increased, there were clear increases in target/standard pixel-wise luminance RMS error and decreases in phase-invariant structural similarity (**Fig. 5, blue lines**). Together, these relationships verify that the target patch is most similar to the standard patch for smaller versus larger separation distances, corroborating previous work (Field, 1987; Frazor & Geisler, 2006; Harrison, 2021; Simoncelli & Olshausen, 2001). In contrast, raw pixel-wise luminance and phase-invariant structural similarity scores for the foil/standard did not vary systematically with separation distance (**Fig. 5, orange**). Further, across all separation distances, the foil was less similar overall to the standard than was the target. This consistently higher similarity of the target than the foil fits well with participants’ high proportion of correct responses across conditions. In sum, we observed systematic effects of the separation manipulations on similarity, verifying the effectiveness of the manipulation of low-level feature correlations via spatial separation.

**Figure 5.**
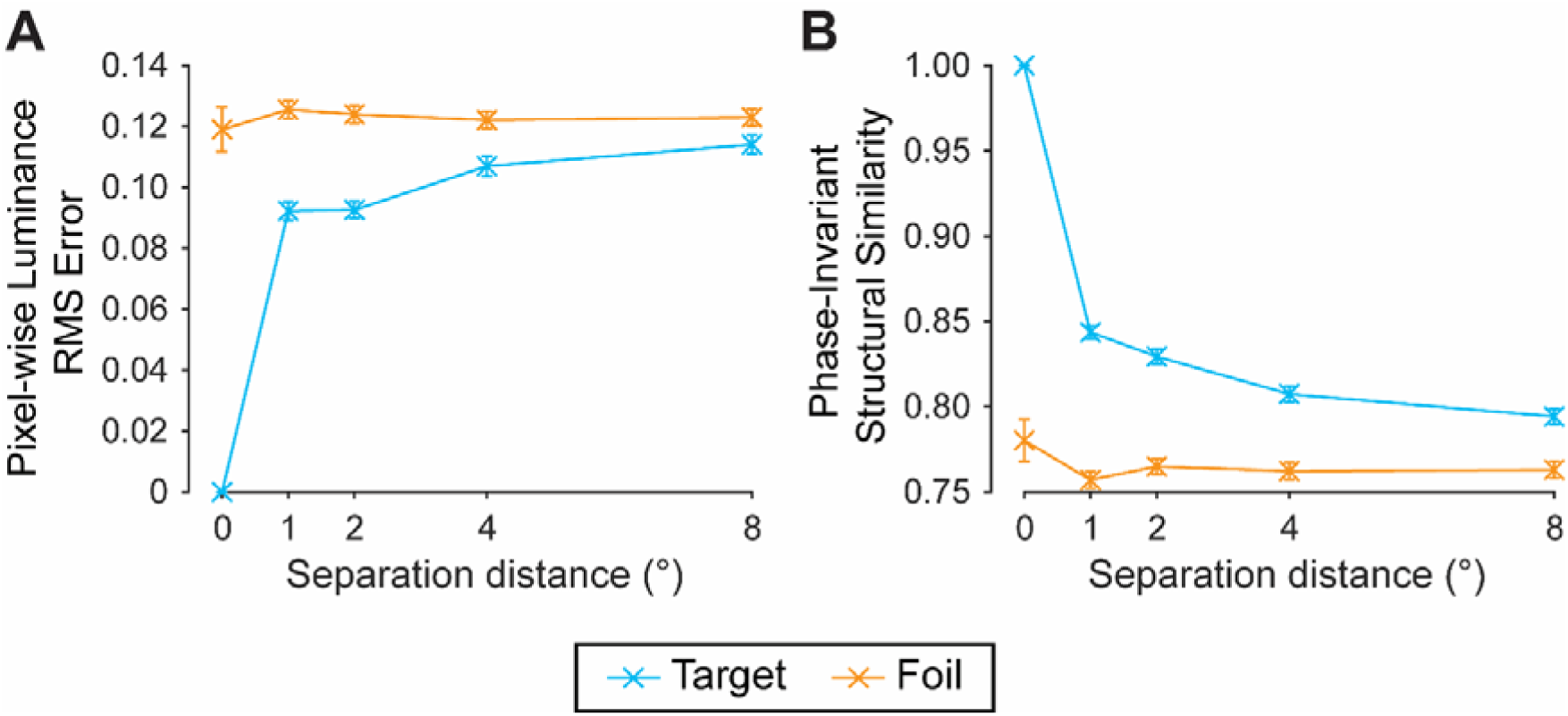
Effect of separation distance on predictor values implemented for the GLMM in Experiment 1. **A)** Mean effect of separation distance on pixel-wise luminance RMS error values, comparing the standard patch with the target (blue) and foil (orange). Note: higher values indicate lower levels of similarity with the standard patch. **B)** Mean effect of separation distance on phase-invariant structural similarity values, comparing the standard patch with the target and foil. Note: higher values indicate higher levels of similarity with the standard patch. Error bars: ±1 SEM (in some cases, standard errors are smaller than the point size). See **S.3** for effect of separation azimuth.

Using predictors based on low-level information, we observed a consistent relationship between the GLMM predictions and participants’ responses (**Fig. 4, solid lines**). Indeed, both pixel-wise luminance and phase-invariant structural similarity were significant predictors in the model (maximum *p* < .001), however the interaction between predictors was not significant (*p* = .983; see **S.4** for full model output). Importantly, the model did not include predictors for separation conditions (i.e., distance and azimuth). Nevertheless, after generating predictions, the data were partialled into separation conditions; they revealed high consistency with behavioural data for each individual condition, including at 0° separation (**Fig. 4**). Overall, the high level of congruence observed between the model and the behavioural data suggests clear explanatory power of low-level features in accounting for participants’ responses.

For the sake of completeness, several other analyses were performed to explore effects of individual separation azimuths, as well as separation fields (e.g., upper vs lower separations relative to horizontal; see **S.5-6**). The GLMM fit all exploratory separation classifications well. Additional inspections will not be discussed in depth because they largely did not reveal clear effects. One exception was the observation of slightly better performance for left separations as compared with right, relative to vertical (see **S.6**). This separation field effect is most likely due to a stimulus set-specific bias that made similarity judgements marginally easier for left separations compared with right separations. The presence of a stimulus bias is supported by the GLMM predictions, which closely corresponded with this response pattern.

### 4.2. Experiment 2: Which low-level features facilitate similarity judgements?

On finding that low-level information correlations can predict participants’ image region associations in Experiment 1, Experiment 2 sought to more directly investigate *which* features facilitate participants’ performance. The same image region association task as in Experiment 1 was implemented, but now the target/foil images were passed through one of three image processing manipulations: “full”, “threshold”, and “edge” processing (**Fig. 3**). All other separation distance and azimuth conditions from Experiment 1 were repeated in Experiment 2, and observers’ responses were again modelled using low-level information predictors.

We explored the impact of separation distance, separation axis, and their interaction on participants’ performance for Experiment 2 for the full image condition to compare patterns of results with those found in Experiment 1 (**Fig. 6**; see **S.7-9** for other image processing conditions). An initial Bayesian 3 (image processing: full, edge, threshold) x 2 (separation axis: cardinal, oblique) x 4 (separation distance: 1°, 2°, 4°, 8°; 0° was not included as this condition could not interact with separation axis) repeated measures ANOVA was conducted and revealed moderate evidence in favour of there being no three-way interaction (BF_10_ = 0.200). Given the lack of three-way interaction and that the purposes of describing behavioural effects is to compare with the patterns observed in Experiment 1, for simplicity, we report the results of a Bayesian 2 (separation axis: cardinal, oblique) x 4 (separation distance: 1, 2, 4, 8) repeated measures ANOVA for the full image condition only.

**Figure 6.**
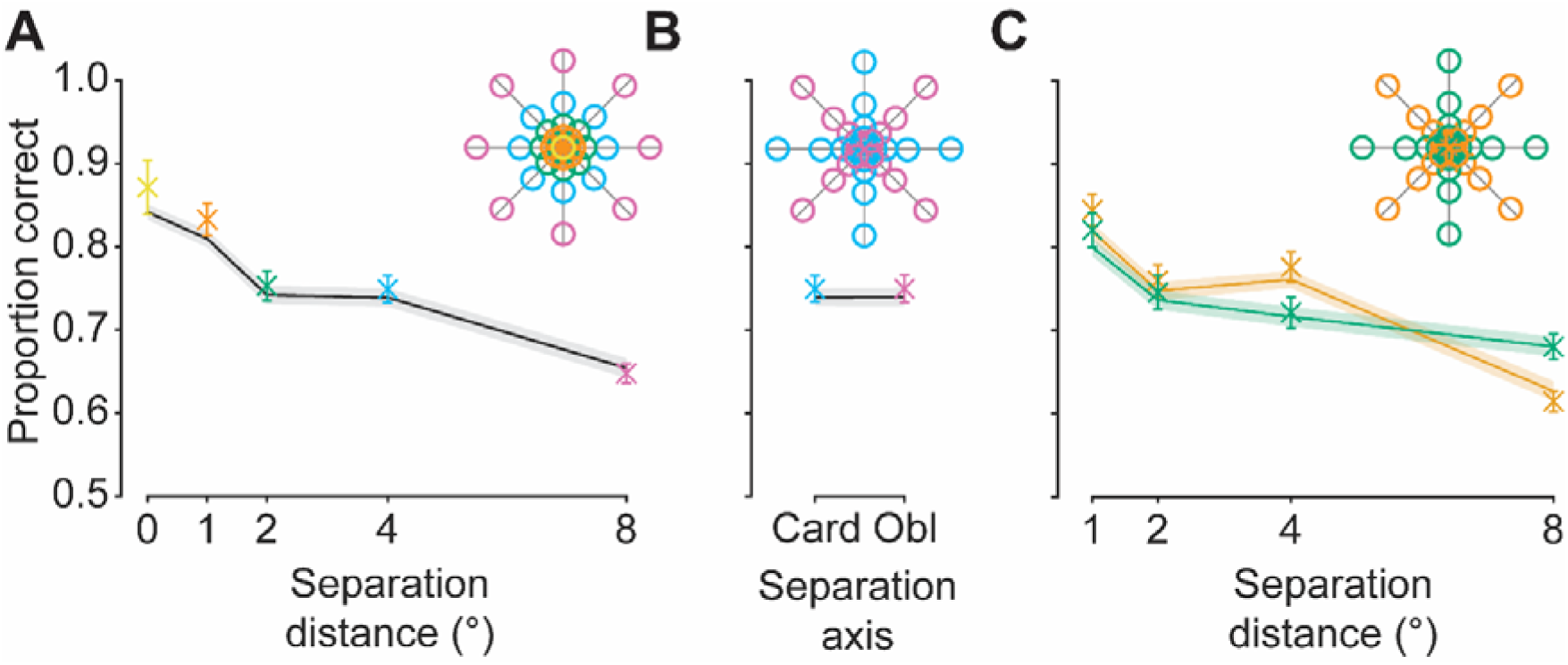
Experiment 2 behavioural and GLMM results for the full image condition only. **A)** Effect of separation distance (x-axis) on the proportion of correct responses (y-axis). Data points are colour coded and are averaged across spatial locations of the same colour in the legend (inset). Solid line represents GLMM predictions based on luminance and structural similarity heuristics between the target and standard patch vs the foil and standard patch. **B)** Effect of separation axis (cardinal vs oblique) on the proportion of correct responses. **C)** Interaction between separation distance (x-axis) and separation axis (separate lines; see legend) on the proportion of correct responses. Error bars: ±1 SEM for participant responses (in some cases, standard errors are smaller than the point size). Shaded regions: ±1 SEM for trial-by-trial model predictions.

As shown in **Figure 6**, increasing separation distance between the standard and the target led to decreased performance in the full image condition, as in Experiment 1, supported by extreme evidence in favour of a separation distance main effect (BF_10_ = 3.295x10^26^). Post-hoc comparisons demonstrated extreme evidence in favour of a difference between all separation distance conditions (minimum BF_10_ = 2.118x10^5^) with the exception of the comparison of the 2° and 4° separation distance conditions, which revealed moderate evidence in favour of there being no difference in performance (BF_10_ = 0.180). The equivocal performance in the 2° and 4° separation distance conditions (see **Fig. 6A**) is likely due to a bias in the stimulus set used for Experiment 2. Specifically, given the random selection of source images, the features participants used to complete the task likely vary between conditions. In this case, it is likely that such features in the 2° and 4° separation distance conditions happened to result in similar difficulty levels. Low accuracy scores were observed for the 0° separation distance condition. We suspect this result, as in Experiment 1, is likely due to a combination of the overall variability in participant-to-participant performance (see **S.7**) and the greater response variance for this datapoint (see **S.10**). There was moderate evidence in favour of no effect of separation axis on performance (BF_10_ = 0.189), with comparable accuracy levels for cardinal and oblique offsets, consistent with the lack of clear separation axis effect found in Experiment 1 (**Fig. 6B**). Extreme evidence was found in favour of an interaction effect between separation axis and separation distance (BF = 1.014x10^4^; **Fig. 6C**), with a very different qualitative appearance of response patterns as compared with Experiment 1 (where there was no evidence in favour of an interaction; comparing **Fig. 4C** with **6C**). This result, as with the dip in performance for the 2° separation distance condition, likely also followed from a bias in the features present in the current stimulus set. The results of Experiment 2, while largely consistent with those of Experiment 1, therefore suggest that the outcomes are influenced by the specific stimulus set used whose features (e.g., low-level pixel-wise information correlations) influence association judgements differentially across separation conditions. We performed several other exploratory analyses, as in Experiment 1, which we do not discuss in depth here (but see **S.7-11** for additional inspections, as well as individual data).

In Experiment 2, participants made similarity judgements on either full, threshold, or edge-only images. Participants performed best in the full patch condition across all separation conditions (**Fig. 7A**; see **S.7-9 & A.11** for individual data and interaction with other separation manipulations), suggesting neither of the image filtering techniques produced stimuli that contained sufficient information to yield the same levels of performance. The initial Bayesian 3 (image processing: full, edge, threshold) x 2 (separation axis: cardinal, oblique) x 4 (separation distance: 1, 2, 4, 8) repeated measures ANOVA revealed extreme evidence in favour of an image processing condition main effect (BF_10_ = 4.135x10^21^). Here, post-hoc comparisons revealed extreme evidence in favour of higher response accuracy in the full image condition as compared with the threshold (BF_10_ = 4.183x10^25^) and edge conditions (BF_10_ = 2.114x10^24^). Further, there was strong evidence in favour of there being no difference in performance between the edge and threshold patch conditions (BF_10_ = 0.088). This was supported by a significant positive correlation between the average response to edge vs threshold versions of each target image (*r^2^* = 0.325, *p* < .001; **Fig. 7B**). These results suggest that the edge and threshold conditions were similarly informative, and that participants potentially used additional features beyond pixel-wise luminance or structural information to perform the task when provided with full images. Alternatively, poorer performance for the threshold and edge image conditions might have followed from low-level information correlations being reduced by the image processing techniques employed.

**Figure 7.**
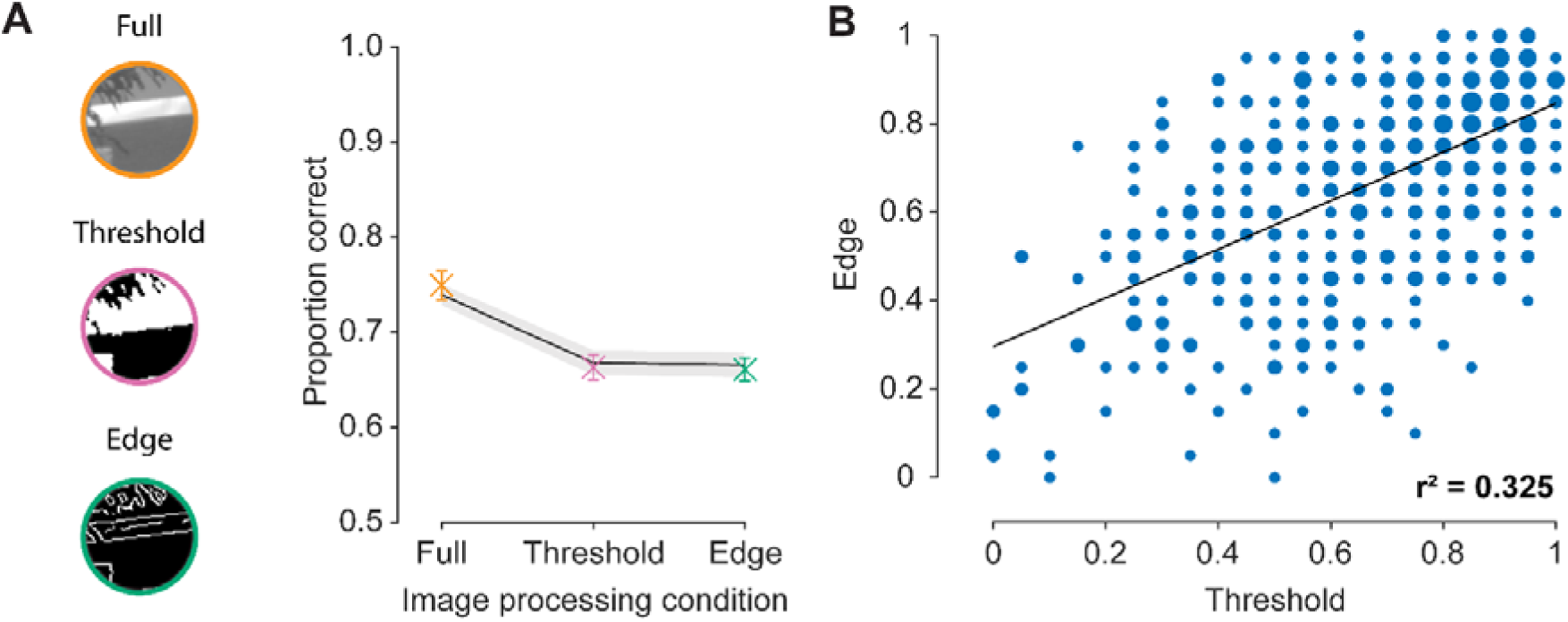
Experiment 2 stimulus examples and results. **A)** Effect of image processing condition (x-axis) on the proportion of correct responses (y-axis). Data points are colour coded according to image conditions depicted to the left of the plot. Solid line represents GLMM predictions based on luminance and structural similarity heuristics between the target and standard patch vs the foil and standard patch. Error bars: ±1 SEM for participant responses. Shaded regions: ±1 SEM for trial-by-trial model predictions. **B)** Correlation between the average response for each threshold image (x-axis) with their corresponding (i.e., generated from the same source image) edge image (y-axis). Values of 1 indicate that the target was always chosen over the foil, and values of 0 indicate the foil was always chosen over the target. Note: there are overlapping data points, indicated by larger symbols.

To investigate the impact of image processing conditions on low-level pixel-wise information correlations, raw pixel-wise luminance and phase-invariant structural similarity scores were inspected. Such scores were computed for patches after having undergone any relevant image processing for a given trial. Indeed, target patches were less similar to the standard after undergoing the edge and threshold processing techniques. Taking pixel-wise luminance as an example, for the full image condition, a similar pattern and similar RMS error values were found to those reported for in Experiment 1 (comparing **Fig. 5A** with **Fig. 8A**). Specifically, as separation distance increased, there was an increase in pixel-wise luminance RMS error (indicating decreasing similarity) for the target, with relatively stable error for the foil across separation distances (**Fig. 8A**). However, we note the similar levels of RMS error between the 2° and 4° conditions for both targets and foils, in line with similar levels of response accuracy. Further, this relationship was much less clear for the threshold and edge image conditions, with far greater overlap in target and foil RMS error scores across separation distance (**Fig. 8B-C**), as well as higher RMS error values overall (comparing **Fig. 8B-C** with **Fig. 8A**). Together, these results indicate that the processing of the patches led to significant disruptions to the pixel information, resulting in less informative low-level information correlations. These lower correlations are consistent with the lower levels of performance observed across the threshold and edge image conditions as compared with the full image condition.

**Figure 8.**
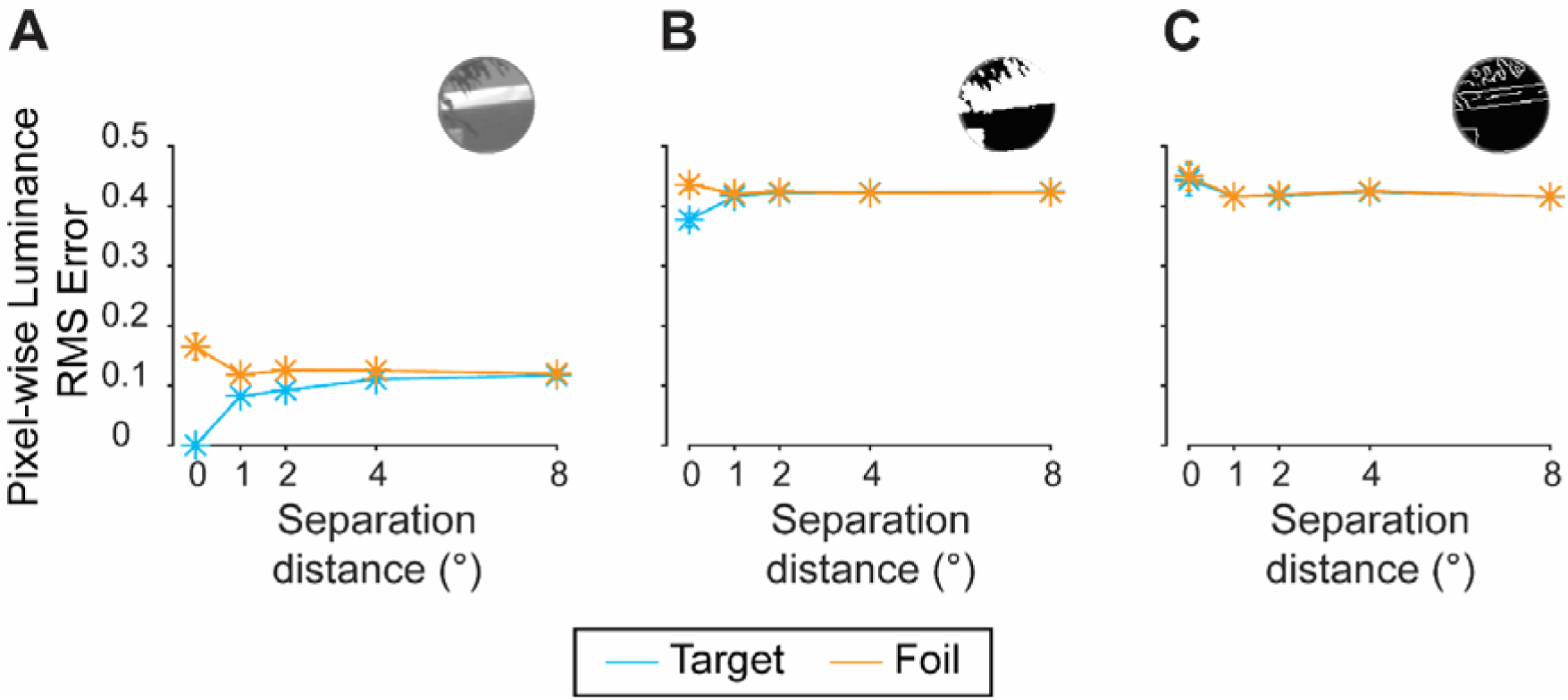
Effect of separation distance on pixel-wise luminance RMS error values, implemented for Experiment 2’s GLMM. **A)** Mean effect of separation distance on pixel-wise luminance RMS error values, comparing the standard patch with the target (blue) and foil (orange) in the *full* image condition. **B)** Mean effect of separation distance on pixel-wise luminance RMS error values, comparing the standard patch with the target and foil in the *threshold* image condition. **C)** Mean effect of separation distance on pixel-wise luminance RMS error values, comparing the standard patch with the target and foil in the *edge* image condition. Error bars: ±1 SEM (in some cases, standard errors are smaller than the point size). See **S.12-13** for overall effect of separation distance, effects of separation azimuth, as well as phase-invariant structural similarity results.

Pixel-wise luminance and phase-invariant structural similarity scores were used as predictors for a GLMM that was fit to Experiment 2 data, as in Experiment 1. As observed in Experiment 1, phase-invariant structural similarity was a significant predictor (*p* = .012) and the interaction between predictors was not significant (*p* = .327). However, unlike in Experiment 1, pixel-wise luminance error was not a significant predictor (*p* = .382), consistent with the larger degree of overlap in raw pixel-wise error scores as a result of image processing (**Fig. 8B-C**), suggesting this predictor was unable to distinguish between the target and foil (see **S.14** for full model output). Nonetheless, across all conditions, there was consistency between model predictions and behavioural data, including at 0° separation.

Model/behavioural data consistency across conditions supports the notion that stimulus set-specific biases can influence low-level feature correlations and participant responses. Specifically, the equivocal performance in the 2° and 4° separation distance conditions, as well as the interaction between separation axis and distance, were both accounted for by GLMM predictions (**Fig. 6**). Further, GLMM predictions were consistent with behavioural performance for each image processing condition, suggesting lower levels of performance for edge and threshold images can also be accounted for by changes in low-level information correlations (**Fig. 7A, solid line**). Hence, the results of Experiment 2 further demonstrate clear explanatory power of low-level features, in particular structural similarity, in accounting for participants’ responses.

## 5. Discussion

We investigated observers’ capacity to associate naturalistic scene regions based on low-level visual features. Across two experiments, participants viewed image patches, windowed from broader naturalistic images, and indicated which two of three were drawn from the same broader scene, allowing us to measure participants’ ability to accurately identify which regions came from the same scene. Critically, we also implemented a GLMM framework to predict responses using low-level (i.e., image computable) information. Specifically, the predictors for the model were pixel-wise luminance and phase-invariant structural similarity. Using a GLMM framework allowed assessment of the potential contribution of such low-level information to image region associations. The GLMM was able to account for participants’ responses, suggesting low-level information is sufficient to facilitate image region associations.

### 5.1. Observers can associate isolated naturalistic image regions

We found evidence that observers are able to associate isolated image regions with one another, building on existing evidence for the computational capacity to perform such associations (Field, 1987; Frazor & Geisler, 2006; Harrison, 2021; Simoncelli & Olshausen, 2001). Importantly, spatial separations between the two patches that *did* belong to the same broader scene were manipulated across experiments – namely, via distance and azimuth manipulations – to introduce systematic variance in low-level feature correlations. In both Experiment 1 and Experiment 2, participants were unaware of the spatial manipulations used, but consistently performed above chance at all separation distances and axes, suggesting our ability to associate regions of space is robust to such spatial offsets.

The results of the current study are consistent with the broader scene perception literature, which has demonstrated that observers can process scene information rapidly and with a high degree of fidelity. For example, observers can perform scene categorisation judgements with presentation times as short as a couple of hundred milliseconds (Fei-Fei et al., 2007; Thorpe et al., 1996; VanRullen & Thorpe, 2001). Unlike past studies that provided participants with “whole” naturalistic images that conveyed entire scenes, however, the current study employed smaller windowed image regions, allowing us to effectively limit the amount of information available in a given image to approximate that of the fovea (A-Izzeddin et al., 2022). For example, in most cases, the windowing of image regions almost completely eliminated the presence of multiple objects that could have indicated the regions’ spatial position within the scene. By providing such limited scene information, the current study expands on previous work, demonstrating that we require very little information to understand properties of the broader scene. Previous research suggests that the presence of such location-based object relationships facilitates perceptual tasks using naturalistic scene stimuli, such as object-based visual search (Brockmole et al., 2006; Brockmole & Henderson, 2006b, 2006a; Eckstein et al., 2006; Loftus & Mackworth, 1978). However, the current study suggests that such relationships at the object-level may not be necessary for scene mapping, with this potentially requiring only very limited visual information.

### 5.2. Judgements on complex stimuli are reducible to low-level feature correlations

Our results suggest that image region associations can be accounted for using low-level feature correlations. We found that the GLMM provides an excellent fit to participant responses using low-level information correlations as predictors. Other potential candidate models could include alternative/additional low-level features, such as contrast and mean luminance. The current model, having provided predictions that are consistent with observed behavioural responses, is a suitable benchmark against which other models can be tested in future. Methods for calculating low-level feature correlations between naturalistic image regions have been established in the literature, demonstrating clear computational capacity for performing such region associations (Field, 1987; Frazor & Geisler, 2006; Harrison, 2021; Simoncelli & Olshausen, 2001). Further, in the contour grouping literature, there is evidence to suggest that contour association judgements largely follow from basic co-occurring image statistics (Elder & Goldberg, 2002; Geisler et al., 2001; Geisler & Perry, 2009). The current study is consistent and extends on this previous computation and contour grouping work, demonstrating that scene region associations are similarly well-accounted for by low-level features.

We have a demonstrated capacity to perceive and make perceptual decisions using low-level features. For example, we know that observers are sensitive to and are able to reproduce individual orientations (Appelle, 1972; Bays, 2014; Berkley et al., 1975; Campbell et al., 1966; Dakin, 2001; de Gardelle et al., 2010; Emsley, 1925; Girshick et al., 2011; Harrison & Bays, 2018; Henderson & Hollingworth, 1999; Pratte et al., 2016; Taylor & Bays, 2018; Westheimer & Beard, 1998) and are even sensitive to the average of an array of orientations (Dakin et al., 2009; Dakin & Watt, 1997; Parkes et al., 2001). Each of these judgements requires the effective processing of available low-level information. The current study extends on this work by investigating perception of such low-level information when it is embedded within more complex stimuli. Indeed, we have found evidence to suggest that judgements on naturalistic image regions can be accounted for by low-level feature correlations alone.

### 5.3. The role of alternative visual information

In Experiment 2, participants were provided with stimuli that were either unaltered, or had undergone processing to reduce information to either thresholded contrast or edge content. Here, perhaps unsurprisingly, participants’ performance dropped substantially when given limited image information in comparison to when given unaltered image regions. Low-level image computable information correlations between the standard/target patches were heavily impacted by the edge and threshold image processing techniques implemented. Specifically, image processing resulted in much lower similarity scores, resembling scores found when comparing the foil/standard patches, suggesting that poorer performance is to be expected if participants are using low-level information. However, there remains the possibility that alternative information was being used to support the higher level of performance observed for unaltered images that is eliminated by further image processing.

A clear candidate for alternative information participants may use in the full image condition is higher-level or semantic information. By higher-level information, we refer to features which convey meaning and are not image computable, such as the arrangement of chairs in a room that denote a living room rather than a dining room (Neri, 2014). A-Izzeddin et al. (2022) had participants infer the upright orientation of randomly oriented image regions. In that study, the same stimulus generation method was employed, cropping the same sized image regions from the same bank of photographs. As part of their study, A-Izzeddin et al. (2022) conducted a control experiment and found that only a very small proportion of stimuli (∼4%) generated using this method conveyed informative high-level information that could disambiguate the patches for the task used. Further, they found that when participants were given patches with informative content, responses were varied, and sometimes yielded large errors in orientation judgements. Hence, even when provided with such informative content, participants do not seem able to effectively utilise such information to inform their judgements. The results of A-Izzeddin et al. (2022) thus suggest that high-level information is unlikely to be a substantial contributor to participants’ performance in the current task when given unaltered image regions. It therefore follows that any further removal of high-level information from the images after undergoing processing is unlikely to explain the observed drop in performance.

Importantly, the GLMM employed in Experiment 1 and Experiment 2 could predict responses using low-level information, across the image processing conditions employed. The GLMM was also able to predict responses across all separation conditions, despite not having access to this spatial information. The ability to predict responses using such low-level information suggests that, even if participants use higher-level visual information to perform the task, we can still account for judgements using the most basic of visual features. There are of course additional visual features that participants might use and which were not accounted for in the current model (e.g., overall patch contrast and mean luminance). Such features, similar to those inspected in the current model, likely interacted with the application of image processing techniques in Experiment 2, which may have driven the poorer performance observed. Future studies could investigate the impact of manipulating other features beyond pixel-wise luminance- and edge-defined structure on image association judgements. Such manipulations would advance our understanding of which features participants actively utilise when performing image region association judgements. However, the performance of the current model suggests that any relevant alternative features follow from, or are highly correlated with, the image computable information used as predictors, leading to our ability to account for participants’ responses to unaltered image regions.

### 5.4. Associating regions of space informing complex scene mapping

Beyond the association of two regions of space, there has been continued speculation surrounding the mechanism by which observers generate a coherent representation of a viewed scene (Berens et al., 2021; Chen et al., 2023; Cohen et al., 1999; Goh et al., 2004; Hassabis & Maguire, 2007; Robertson et al., 2016; Singer & Gray, 1995; Steel et al., 2021; Treisman, 1998). The efficiency with which we process scene information necessitates we be highly adept at this process to enable coherent representations of the scene we are viewing on behaviourally-relevant timescales. Further work is needed to more directly address the specific mechanisms underlying broader scene mapping, as well as the features most “useful” in making such associations. The current study takes a modest step in this direction by showing that we are indeed able to make prompted scene association judgements based on limited available information. In particular, we found that these associations can be reduced to low-level feature correlations, providing evidence for the contribution of such basic visual information to judgements made on complex stimuli.

## Data availability

Behavioural data and GLMM code are available on the Open Science Framework: https://osf.io/ykerd/?view_only=0d3bbfc76768449eb59d3ad011d51979

## Acknowledgements

We would like to thank Laura Wang for her assistance with data collection. EJA was supported by the Deutsche Forschungsgemeinschaft (DFG, German Research Foundation) – SFB/TRR 135 (project no. 222641018, project C1). JBM was supported by a National Health and Medical Research Council (NHMRC; Australia) Investigator Grant (GNT2010141). WJH was supported by an Australian Research Council Discovery Early Career Researcher Award (DE190100136).

## Competing interests

The authors have no conflicts of interest to declare.

## Notes

https://osf.io/ykerd/?view_only=0d3bbfc76768449eb59d3ad011d51979

### Competing Interest Statement

The authors have declared no competing interest.

