## Supplemental Materials for "Low-level features predict perceived similarity for naturalistic images"

S.1. Experiment 1 individual data

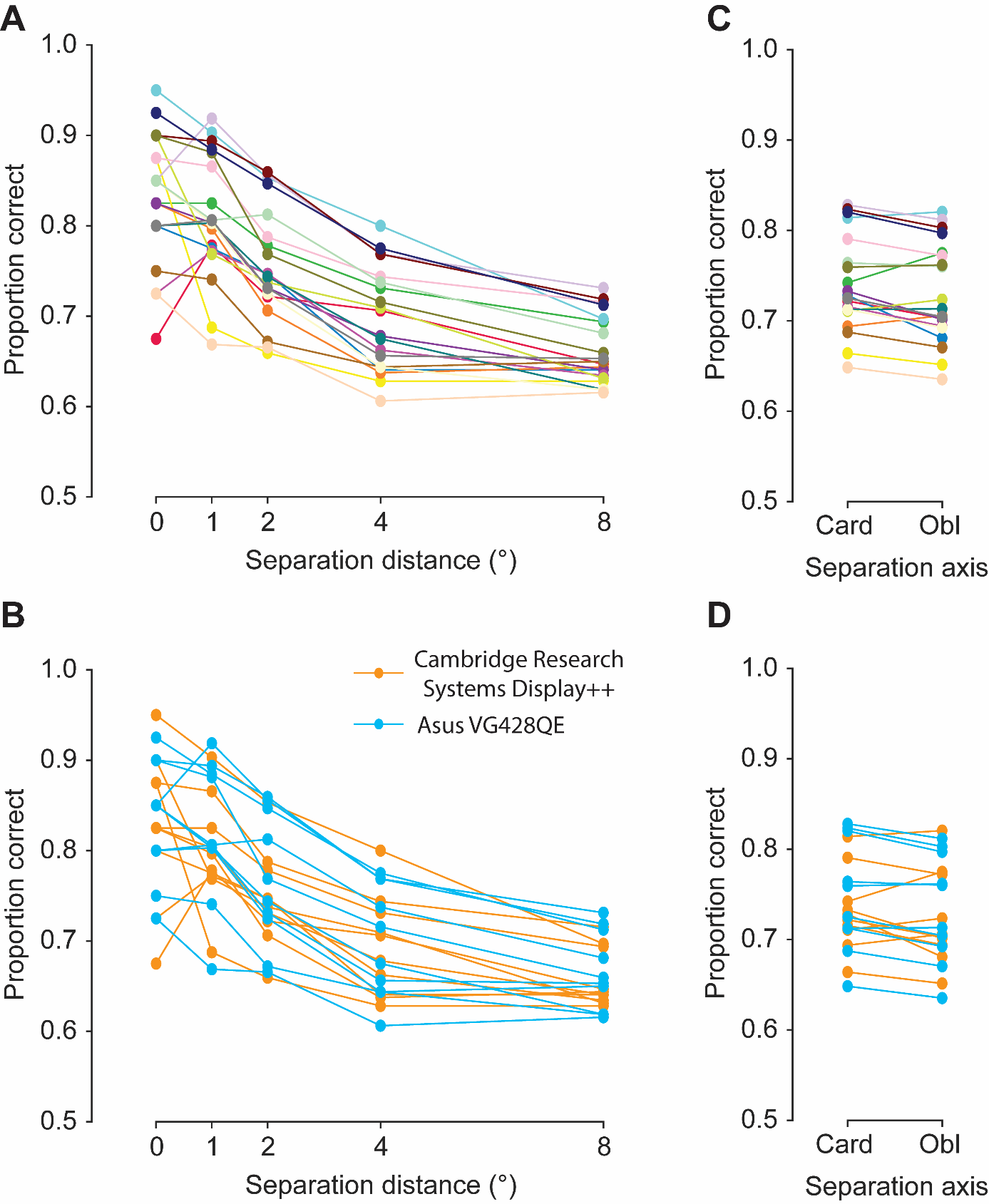

Figure S.1. Individual data for Experiment 1. A) Individual data corresponding to Figure 4A, showing the effect of separation distance (x-axis) on the proportion of correct responses (y-axis). B) The same data as in Panel A, but colour-coded to show participants who participated with the two different monitor set-ups (see inset legend) as outlined in Section 3.4., Apparatus. We see no consistent effect of monitor. C) Individual data corresponding to Figure 4B, showing the effect of separation axis (cardinal vs oblique) on the proportion of correct responses. D) The same data as in Panel C, but colour-coded to show participants who participated with the two different monitor set-ups (see legend) as outlined in Section 3.4., Apparatus. We see no consistent effect of monitor. Solid lines are used to connect individual participants’ datapoints and do not represent model fits.

S.2. Experiment 1 variance data

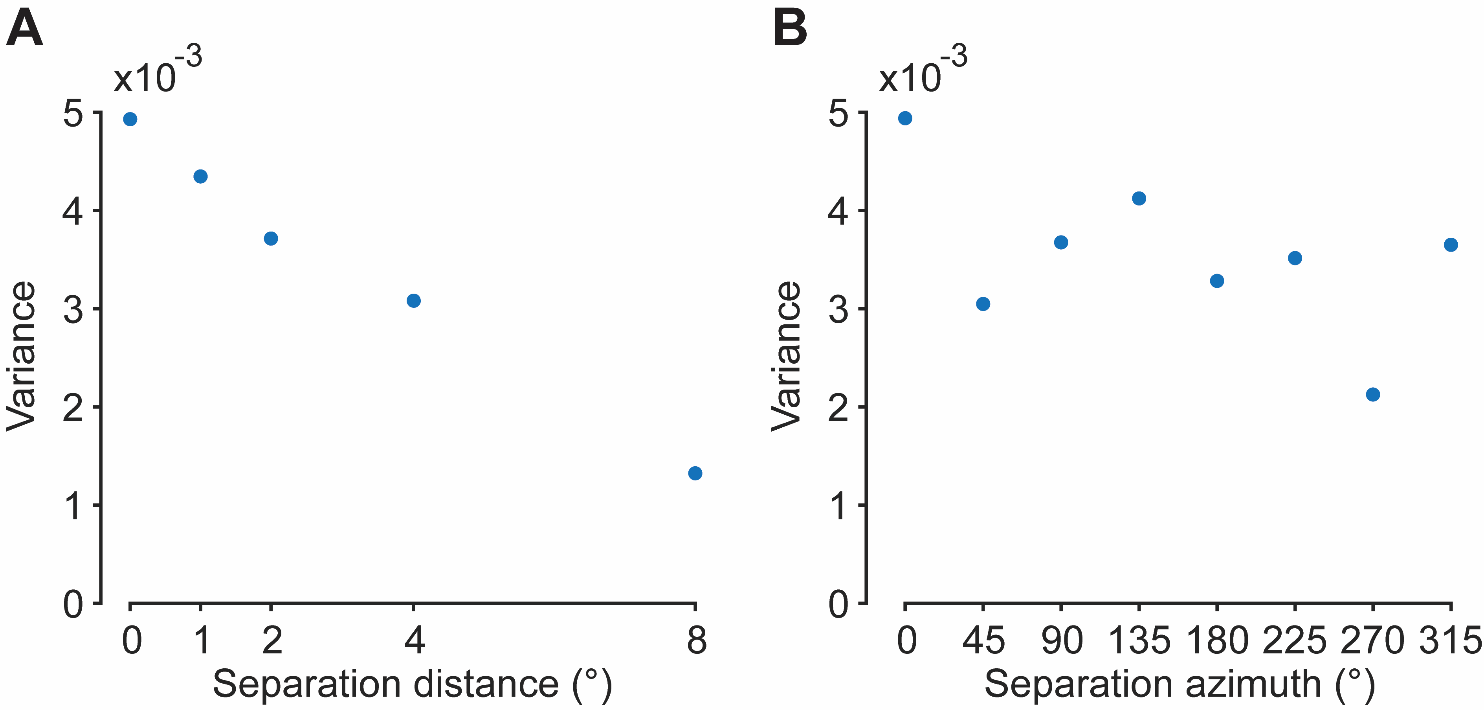

Figure S.2. Experiment 1 variance data corresponding to response accuracy data presented in Figure 4A and Figure S.4A. The effect of separation distance (A; x-axis) and azimuth (B; x-axis) on response variance (y-axis). Variance was calculated using MATLAB’s var() function, finding the variance in participants’ proportion of correct responses at each separation distance/azimuth condition.

S.3. Effect of separation azimuth on Experiment 1 GLMM predictor values

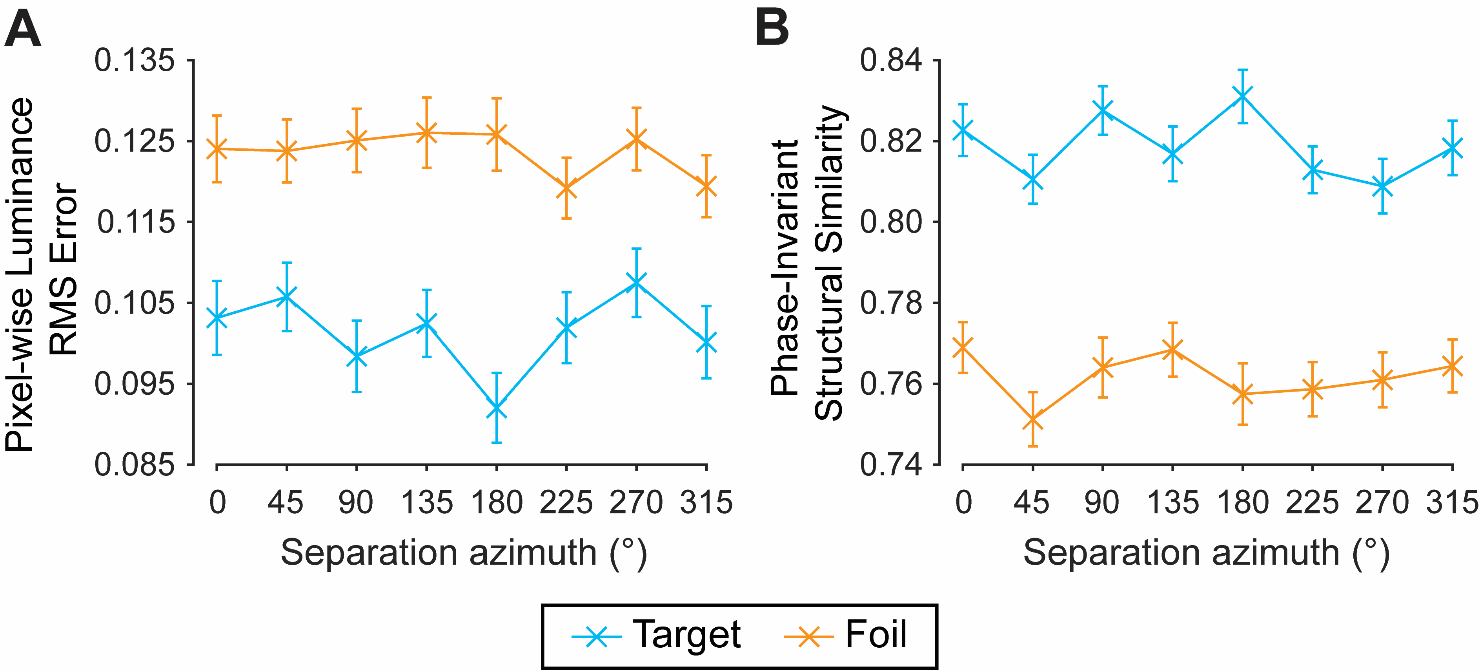

Figure S.3. Effect of separation azimuth on predictor values implemented for Experiment 1’s GLMM. A) Mean effect of separation azimuth on pixel-wise luminance RMS error values, comparing the standard patch with the target (blue) and foil (orange). B) Mean effect of separation azimuth on phase-invariant structural similarity values, comparing the standard patch with the target and foil. Error bars: ±1 SEM.

S.4. Full Experiment 1 GLMM output and comparison

Table S.1: Full output for the Experiment 1 GLMM defined by the equation: $\boldsymbol{y \sim}\boldsymbol{\beta}_{\boldsymbol{0}}\boldsymbol{+}\boldsymbol{\beta}_{\boldsymbol{1}}\boldsymbol{I}_{\boldsymbol{\Delta}}\boldsymbol{+}\boldsymbol{\beta}_{\boldsymbol{2}}\boldsymbol{S}_{\boldsymbol{\Delta}}\boldsymbol{+}\boldsymbol{\beta}_{\boldsymbol{3}}\boldsymbol{I}_{\boldsymbol{\Delta}}\boldsymbol{S}_{\boldsymbol{\Delta}}$. Here, $\boldsymbol{\beta}_{\boldsymbol{0}}$ is the intercept term, $\boldsymbol{\beta}_{\boldsymbol{1}}$ is the weight of the pixel-wise luminance difference, $\boldsymbol{I}_{\boldsymbol{\Delta}}$, $\boldsymbol{\beta}_{\boldsymbol{2}}$ is the weight of phase-invariant structural similarity, $\boldsymbol{S}_{\boldsymbol{\Delta}}$, and $\boldsymbol{\beta}_{\boldsymbol{3}}$ is the weight of the interaction $\boldsymbol{I}_{\boldsymbol{\Delta}}\boldsymbol{S}_{\boldsymbol{\Delta}}$. To partially pool coefficient estimates across participants, the GLMM included participant and image combination as random effects.

| **Name** | **Estimate** | **SE** | **tStat** | **DF** | **pValue** |
| --- | --- | --- | --- | --- | --- |
| Intercept | 1.100 | 0.079 | 13.853 | 26396 | <.001 |
| $S_{\Delta}$ | -0.129 | 0.032 | -4.003 | 26396 | <.001 |
| $I_{\Delta}$ | 0.217 | 0.033 | 6.526 | 26396 | <.001 |
| $I_{\Delta}S_{\Delta}$ | -0.001 | 0.029 | -0.021 | 26396 | .983 |

Table S.2: Full output for the Experiment 1 alternative GLMM defined by the equation: $\boldsymbol{y \sim}\boldsymbol{\beta}_{\boldsymbol{0}}\boldsymbol{+}\boldsymbol{\beta}_{\boldsymbol{1}}\boldsymbol{S}_{\boldsymbol{\Delta}}$. Here, $\boldsymbol{\beta}_{\boldsymbol{0}}$ is the intercept term, $\boldsymbol{\beta}_{\boldsymbol{1}}$ is the weight of the phase-invariant structural similarity, $\boldsymbol{S}_{\boldsymbol{\Delta}}$. To partially pool coefficient estimates across participants, the GLMM included participant and image combination as random effects.

| **Name** | **Estimate** | **SE** | **tStat** | **DF** | **pValue** |
| --- | --- | --- | --- | --- | --- |
| Intercept | 1.139 | 0.079 | 14.342 | 26398 | <.001 |
| $S_{\Delta}$ | -0.151 | 0.033 | -4.619 | 26398 | <.001 |

Table S.3: Full output for the Experiment 1 alternative GLMM defined by the equation: $\boldsymbol{y \sim}\boldsymbol{\beta}_{\boldsymbol{0}}\boldsymbol{+}\boldsymbol{\beta}_{\boldsymbol{1}}\boldsymbol{I}_{\boldsymbol{\Delta}}\boldsymbol{+}\boldsymbol{\beta}_{\boldsymbol{2}}\boldsymbol{S}_{\boldsymbol{\Delta}}$. Here, $\boldsymbol{\beta}_{\boldsymbol{0}}$ is the intercept term, $\boldsymbol{\beta}_{\boldsymbol{1}}$ is the weight of the pixel-wise luminance difference, $\boldsymbol{I}_{\boldsymbol{\Delta}}$, $\boldsymbol{\beta}_{\boldsymbol{2}}$ is the weight of phase-invariant structural similarity, $\boldsymbol{S}_{\boldsymbol{\Delta}}$, and $\boldsymbol{\beta}_{\boldsymbol{3}}$ is the weight of the interaction $\boldsymbol{I}_{\boldsymbol{\Delta}}\boldsymbol{S}_{\boldsymbol{\Delta}}$. To partially pool coefficient estimates across participants, the GLMM included participant and image combination as random effects.

| **Name** | **Estimate** | **SE** | **tStat** | **DF** | **pValue** |
| --- | --- | --- | --- | --- | --- |
| Intercept | 1.100 | 0.079 | 13.866 | 26397 | <.001 |
| $S_{\Delta}$ | -0.129 | 0.032 | -4.004 | 26397 | <.001 |
| $I_{\Delta}$ | 0.217 | 0.032 | 6.733 | 26397 | <.001 |

Table S.4: Formal model comparison. Here, we compare the three models described above, corresponding to the structural similarity-only model (S), the main effect model (M) including both structural similarity and pixel-wise luminance difference, and interaction model (I). Absolute ΔAIC, likelihood ratio statistics (LRStat), and *p* values are provided, calculated relative to the winning model (S).

| **Model** | **DF** | **ΔAIC** | **LRStat** | **pValue** |
| --- | --- | --- | --- | --- |
| S | 4 | 0 |  |  |
| M | 5 | 1.472 | 0.528 | .467 |
| I | 6 | 3.527 | 0.473 | .790 |

S.5. Effect of separation azimuth on Experiment 1 behavioural responses

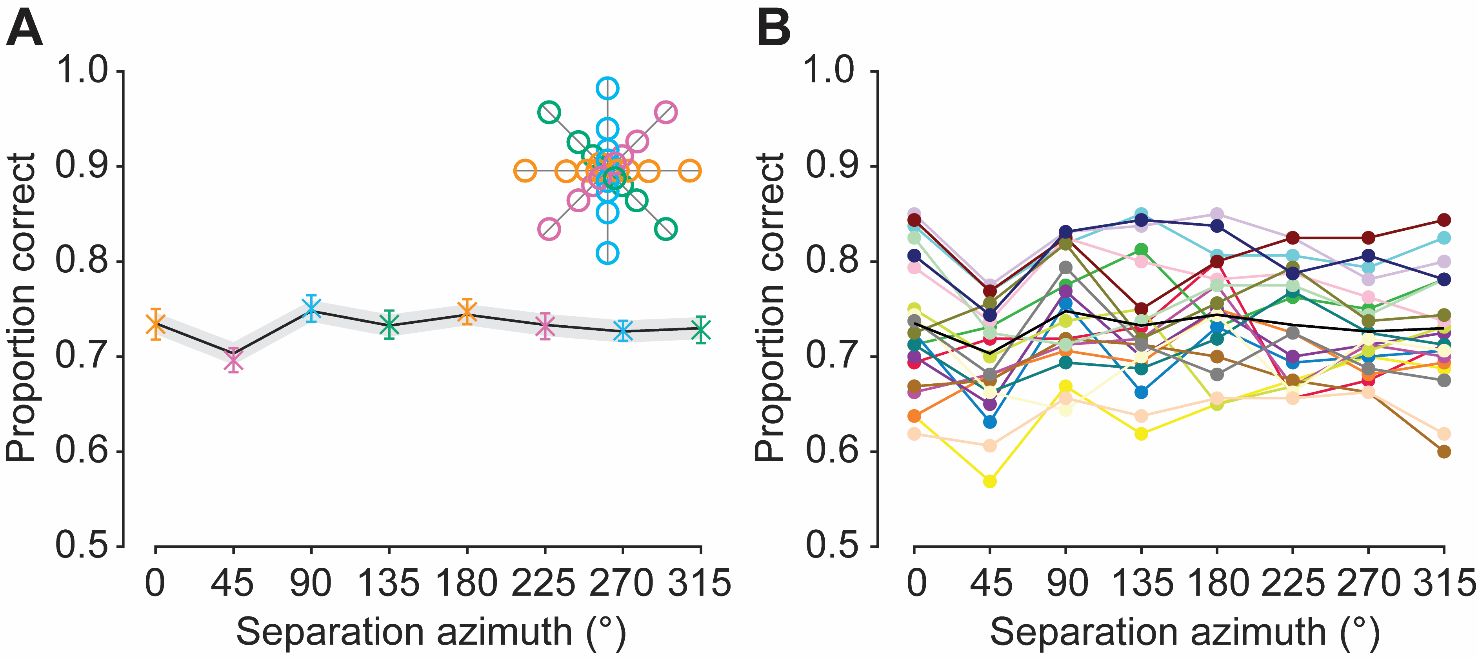

Figure S.4. Separation azimuth results for Experiment 1. A) Effect of separation azimuth on the proportion of correct responses. Data points are colour coded and are averaged across spatial locations of the same colour in the legend (inset). Solid line represents the fits of the generalised linear multilevel model outlined in Section 3.7.2, Generalised linear multilevel modelling. Error bars: ±1 SEM for participant responses (in some cases, standard errors are smaller than the point size). Shaded regions: ±1 SEM for trial-by-trial model predictions. B) Individual data showing the effect of separation azimuth on the proportion of correct responses. Solid lines are used to connect individual participants’ datapoints and do not represent model fits.

S.6. Experiment 1 separation field results

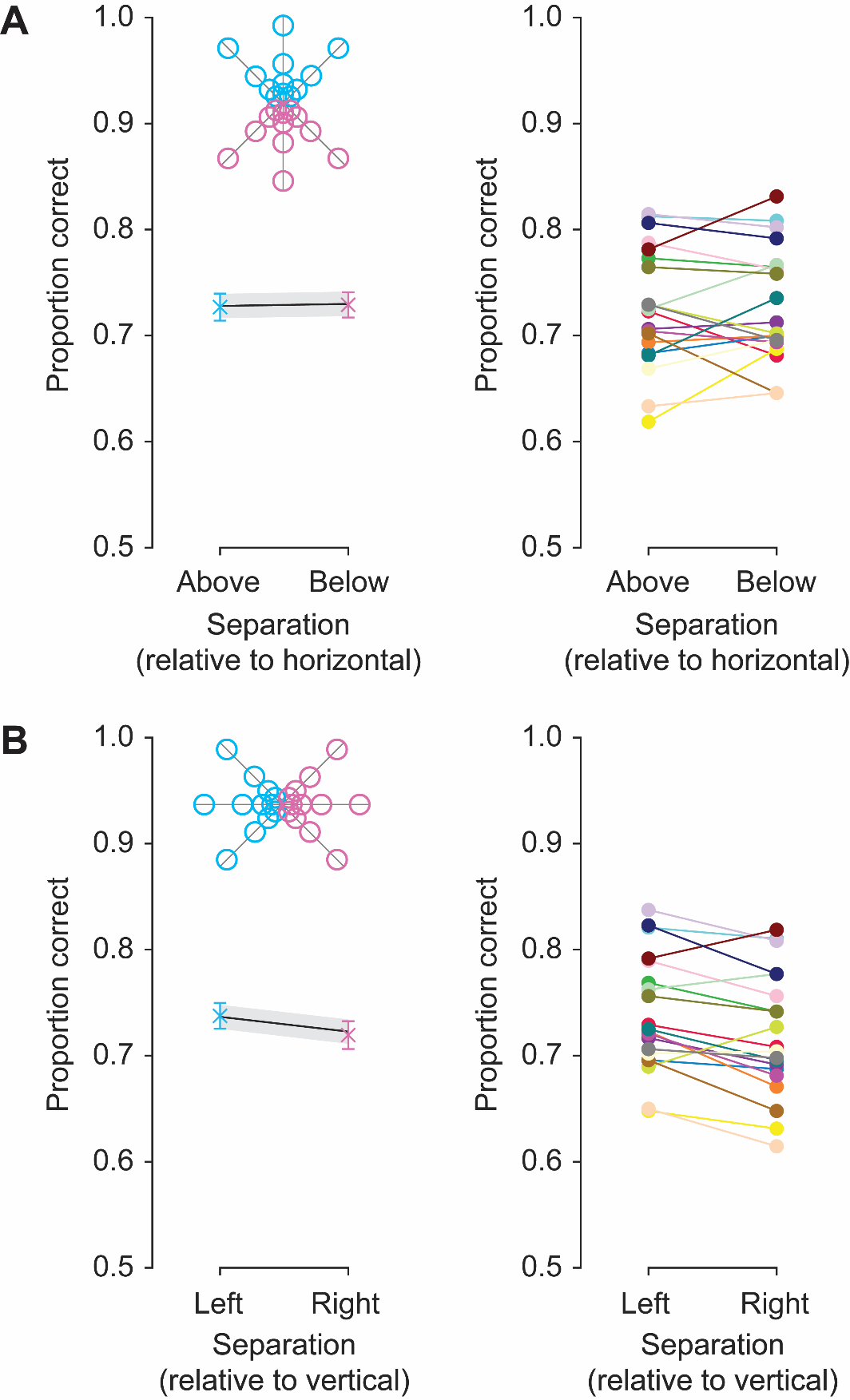

Figure S.5. Separation field results for Experiment 1. A) Effect of separation field relative to horizontal (above vs below) on the proportion of correct responses, with average (left) and individual (right) data in separate plots. Data points are colour coded and are averaged across spatial locations of the same colour in the legend above. B) Effect of separation field relative to vertical (left vs right) on the proportion of correct responses, with average and individual data on separate plots. Data points are colour coded and are averaged across spatial locations of the same colour in the legend above. Error bars: ±1 SEM for participant responses. Solid lines on the left plots represent the fits of the generalised linear multilevel model outlined in Section 3.7.2, Generalised linear multilevel modelling. Shaded regions: ±1 SEM for trial-by-trial model predictions. Solid lines on the right plots are used to connect individual participants’ datapoints and do not represent model fits.

S.7. Effect of separation distance and azimuth on Experiment 2 behavioural responses

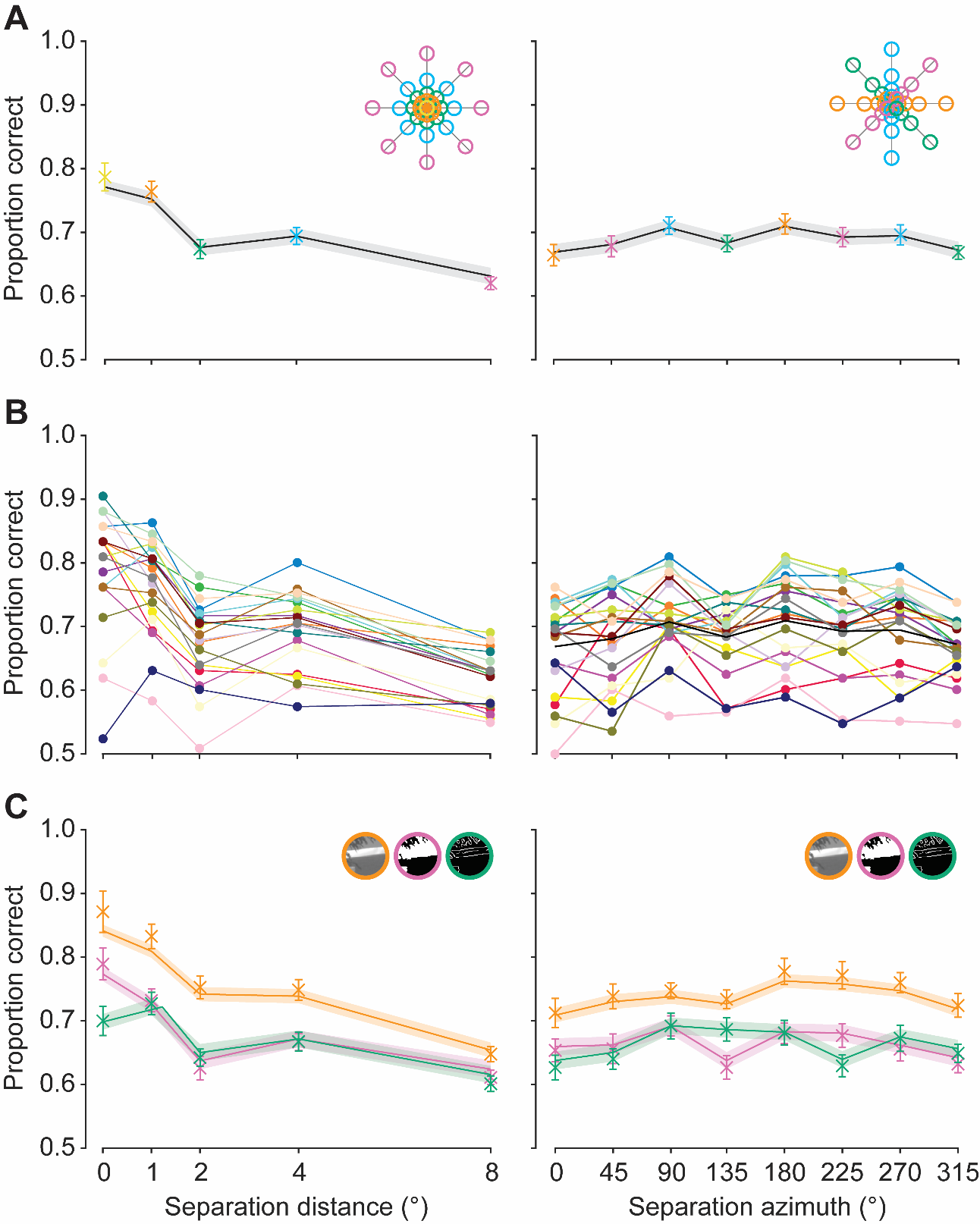

Figure S6. Separation distance (left column) and azimuth (right column) results for Experiment 2. A) Mean effect of distance (left) and azimuth (right) on the proportion of correct responses across participants. Data points are colour coded and are averaged across spatial locations of the same colour in the legend (inset, top right). Solid lines represent the fits of the generalised linear multilevel model outlined in Section 3.7.2, Generalised linear multilevel modelling. Shaded regions: ±1 SEM for trial-by-trial model predictions. B) Individual data, showing the overall effect of separation distance (left) and azimuth (right) on the proportion of correct responses. Solid lines are used to connect individual participants’ datapoints and do not represent model fits. C) Interaction between separation distance (left)/azimuth (right) and image processing condition (separate lines, see top-right inset legend). Solid lines represent the fits of the generalised linear multilevel model outlined in Section 3.7.2, Generalised linear multilevel modelling. Shaded regions: ±1 SEM for trial-by-trial model predictions. Error bars: ±1 SEM for participant responses (in some cases, standard errors are smaller than the point size).

S.8. Experiment 2 separation field results

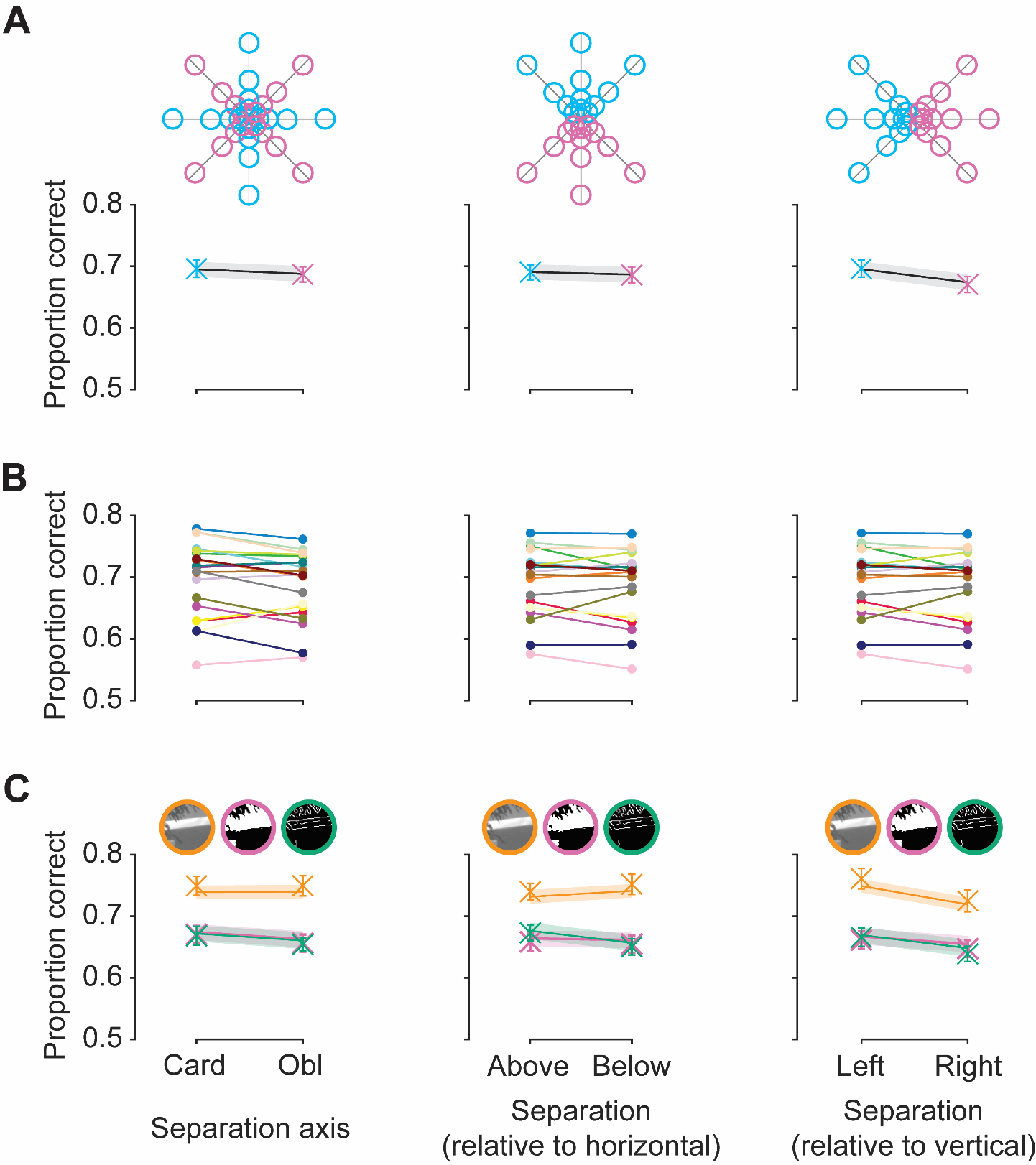

Figure S.7. Separation axis (cardinal vs oblique, left column) and field (upper vs lower, middle column; left vs right, right column) results for Experiment 2. A) Mean effect of separation axis/fields on the proportion of correct responses across participants. Data points are colour coded and are averaged across spatial locations of the same colour in the legend (inset, above). Solid lines represent the fits of the generalised linear multilevel model outlined in Section 3.7.2, Generalised linear multilevel modelling. Shaded regions: ±1 SEM for trial-by-trial model predictions. B) Individual data, showing the overall effect of v axis and fields on the proportion of correct responses. Solid lines are used to connect individual participants’ datapoints and do not represent model fits. C) Interaction between separation axis/fields and image processing condition (separate lines, see above inset legend). Solid lines represent the fits of the generalised linear multilevel model outlined in Section 3.7.2, Generalised linear multilevel modelling. Shaded regions: ±1 SEM for trial-by-trial model predictions. Error bars: ±1 SEM for participant responses (in some cases, standard errors are smaller than the point size).

S.9. Experiment 2 separation distance/axis interaction

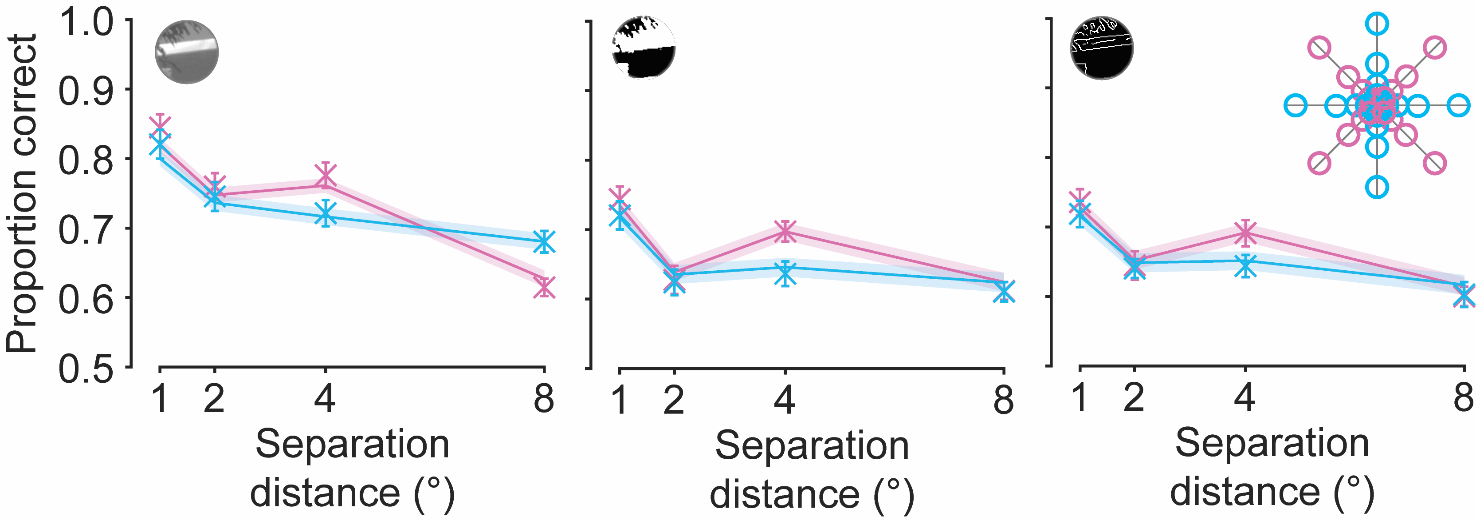

Figure S.8. Separation distance/axis interaction for Experiment 2 for different image processing conditions. Separate separation distance (x-axis) and separation axis (separate lines; see legend inset) interaction results, for each individual image processing condition (as indicated by the top left image in each plot). Solid lines represent the fits of the generalised linear multilevel model outlined in Section 3.7.2, Generalised linear multilevel modelling. Error bars: ±1 SEM for participant responses (in some cases, standard errors are smaller than the point size). Shaded regions: ±1 SEM for trial-by-trial model predictions.

S.10. Experiment 2 variance data

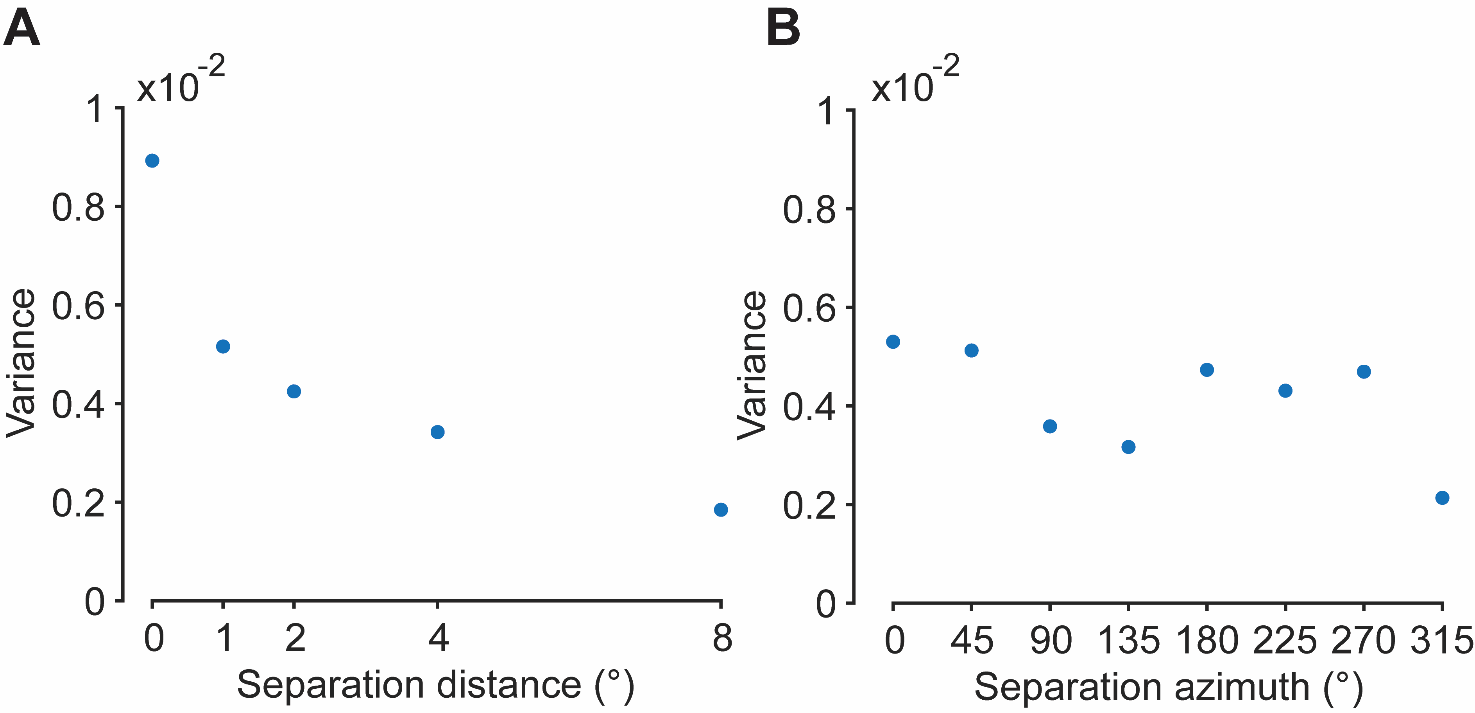

Figure S.9. Experiment 2 variance data corresponding to response accuracy data presented in Figure 6A and Figure S.6A. Plots show the effect of separation distance (A; x-axis) and azimuth (B; x-axis) on response variance (y-axis). Variance was calculated using MATLAB’s var() function.

S.11. Experiment 2 individual data across image processing conditions

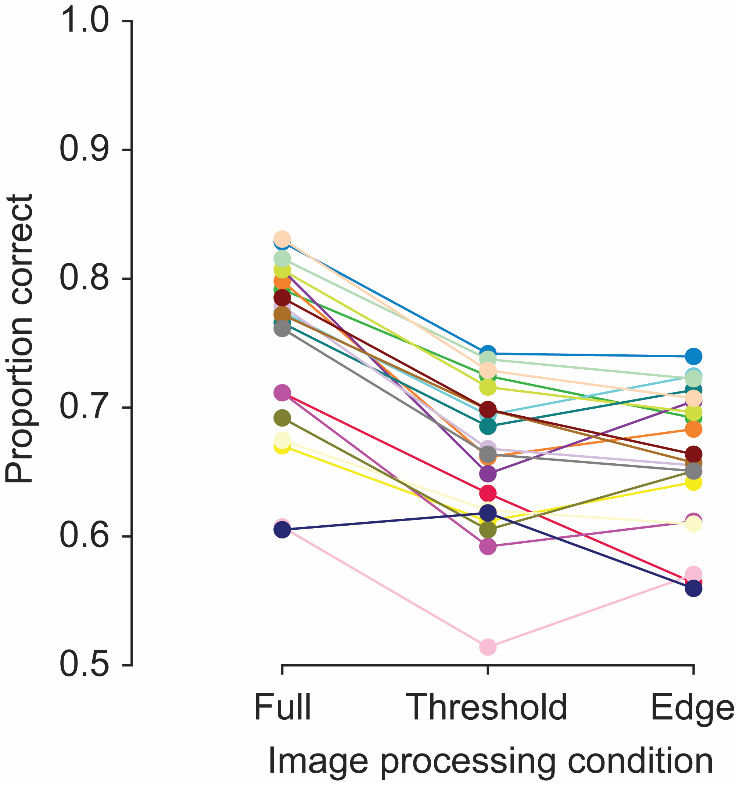

Figure S.10. Individual data corresponding to Figure 7A. The effect of image processing condition (x-axis) is plotted against the proportion of correct responses (y-axis). Solid lines are used to connect individual participants’ datapoints and do not represent model fits.

S.12. Effect of separation on pixel-wise luminance error values for Experiment 2

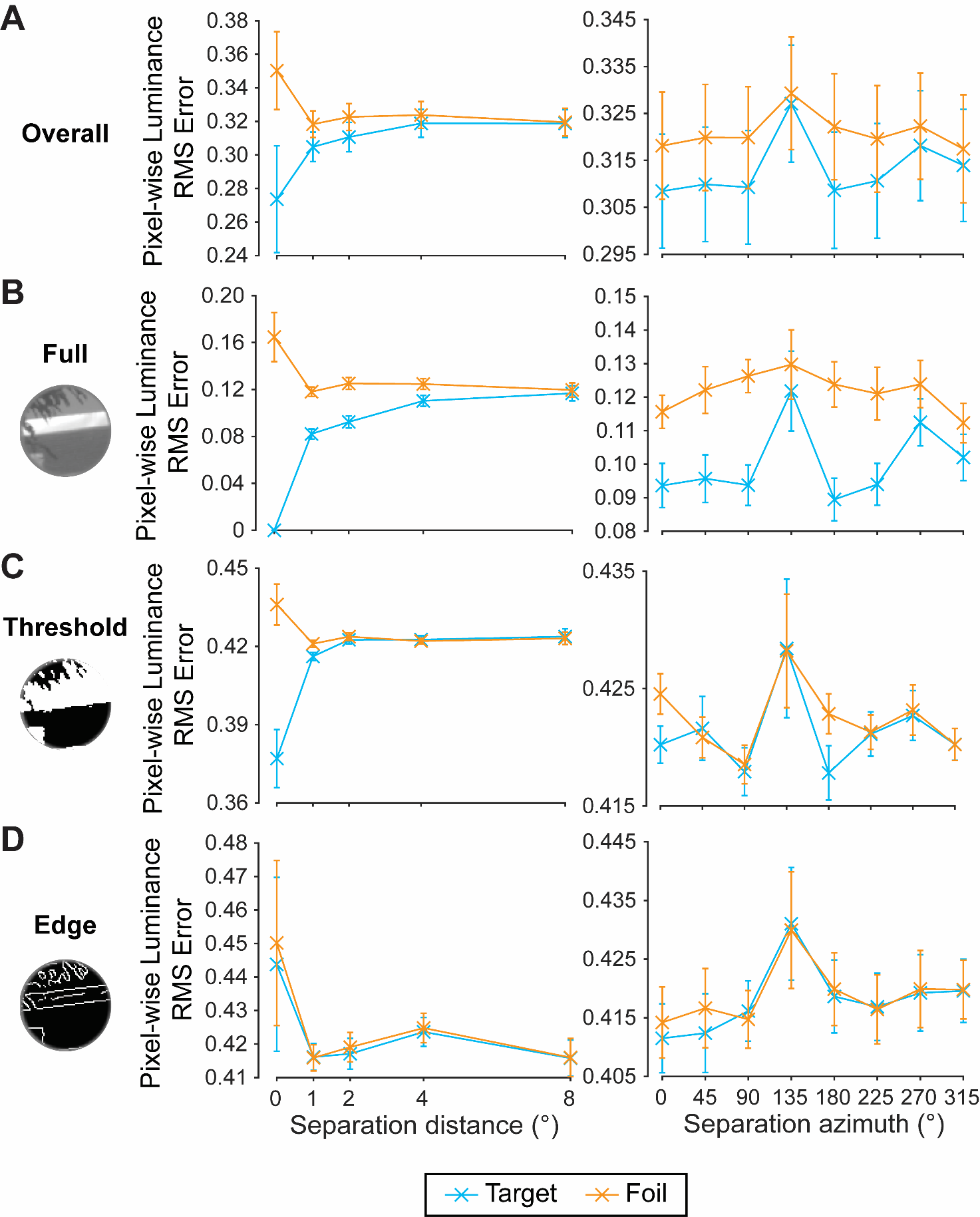

Figure S.11. Effect of separation conditions on pixel-wise luminance error values implemented for the GLMM in Experiment 2. A) Overall effect of separation distance (left) and azimuth (right) on pixel-wise luminance error values, comparing the standard patch with the target (blue) and foil (orange). B-D) Mean effect of separation distance and azimuth on pixel-wise luminance error values, for full, threshold, and edge image processing conditions, respectively. Each plot compares the standard patch with the target and foil patches individually. Error bars: ±1 SEM (in some cases, standard errors are smaller than the point size).

S.13. Effect of separation on phase-invariant structural similarity values for Experiment 2

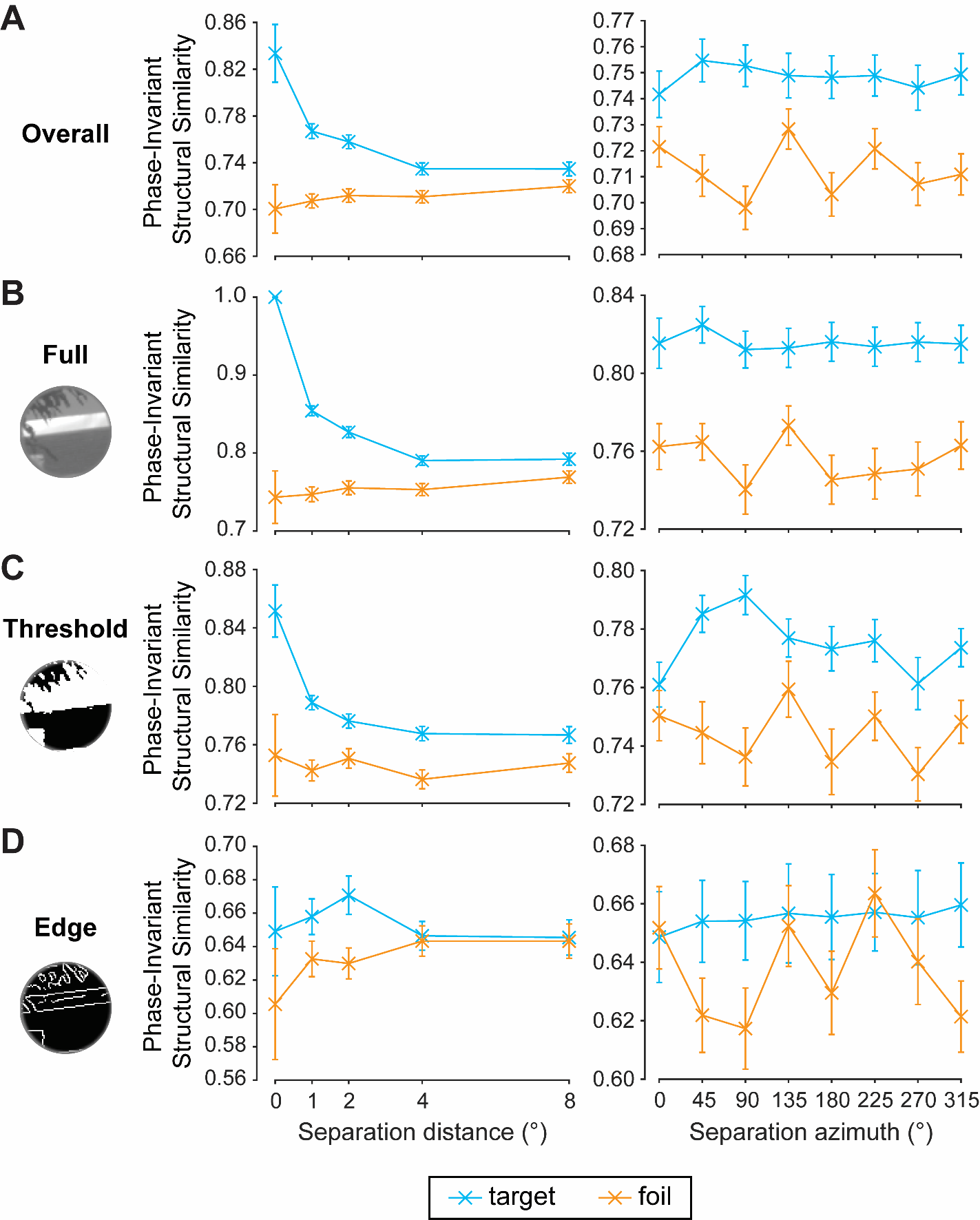

Figure S.12. Effect of separation conditions on phase-invariant structural similarity values implemented for the GLMM in Experiment 2. A) Overall effect of separation distance (left) and azimuth (right) on phase-invariant structural similarity values, comparing the standard patch with the target (blue) and foil (orange). B-D) Mean effect of separation distance and azimuth on phase-invariant structural similarity values, for full, threshold, and edge image processing conditions, respectively. Each plot compares the standard patch with the target and foil patches individually. Error bars: ±1 SEM (in some cases, standard errors are smaller than the point size).

S.14. Full Experiment 2 GLMM output

Table S.5: Full output for the Experiment 2 GLMM defined by the equation: $\boldsymbol{y \sim}\boldsymbol{\beta}_{\boldsymbol{0}}\boldsymbol{+}\boldsymbol{\beta}_{\boldsymbol{1}}\boldsymbol{I}_{\boldsymbol{\Delta}}\boldsymbol{+}\boldsymbol{\beta}_{\boldsymbol{2}}\boldsymbol{S}_{\boldsymbol{\Delta}}\boldsymbol{+}\boldsymbol{\beta}_{\boldsymbol{3}}\boldsymbol{I}_{\boldsymbol{\Delta}}\boldsymbol{S}_{\boldsymbol{\Delta}}$. Here, $\boldsymbol{\beta}_{\boldsymbol{0}}$ is the intercept term, $\boldsymbol{\beta}_{\boldsymbol{1}}$ is the weight of the pixel-wise luminance difference, $\boldsymbol{I}_{\boldsymbol{\Delta}}$, $\boldsymbol{\beta}_{\boldsymbol{2}}$ is the weight of phase-invariant structural similarity, $\boldsymbol{S}_{\boldsymbol{\Delta}}$, and $\boldsymbol{\beta}_{\boldsymbol{3}}$ is the weight of the interaction $\boldsymbol{I}_{\boldsymbol{\Delta}}\boldsymbol{S}_{\boldsymbol{\Delta}}$. To partially pool coefficient estimates across participants, the GLMM included participant and image combination as random effects.

| **Name** | **Estimate** | **SE** | **tStat** | **DF** | **pValue** |
| --- | --- | --- | --- | --- | --- |
| Intercept | 0.966 | 0.079 | 12.239 | 27656 | <.001 |
| $S_{\Delta}$ | -0.088 | 0.035 | -2.525 | 27656 | 0.012 |
| $I_{\Delta}$ | -0.031 | 0.035 | -0.874 | 27656 | 0.382 |
| $I_{\Delta}S_{\Delta}$ | 0.032 | 0.032 | 0.981 | 27656 | 0.327 |

Table S.6: Full output for the Experiment 2 alternative GLMM defined by the equation: $\boldsymbol{y \sim}\boldsymbol{\beta}_{\boldsymbol{0}}\boldsymbol{+}\boldsymbol{\beta}_{\boldsymbol{1}}\boldsymbol{S}_{\boldsymbol{\Delta}}$. Here, $\boldsymbol{\beta}_{\boldsymbol{0}}$ is the intercept term, $\boldsymbol{\beta}_{\boldsymbol{1}}$ is the weight of the phase-invariant structural similarity, $\boldsymbol{S}_{\boldsymbol{\Delta}}$. To partially pool coefficient estimates across participants, the GLMM included participant and image combination as random effects.

| **Name** | **Estimate** | **SE** | **tStat** | **DF** | **pValue** |
| --- | --- | --- | --- | --- | --- |
| Intercept | 0.960 | 0.079 | 12.181 | 27658 | <.001 |
| $S_{\Delta}$ | -0.084 | 0.035 | -2.415 | 27658 | 0.016 |

Table S.7: Full output for the Experiment 2 alternative GLMM defined by the equation: $\boldsymbol{y \sim}\boldsymbol{\beta}_{\boldsymbol{0}}\boldsymbol{+}\boldsymbol{\beta}_{\boldsymbol{1}}\boldsymbol{I}_{\boldsymbol{\Delta}}\boldsymbol{+}\boldsymbol{\beta}_{\boldsymbol{2}}\boldsymbol{S}_{\boldsymbol{\Delta}}$. Here, $\boldsymbol{\beta}_{\boldsymbol{0}}$ is the intercept term, $\boldsymbol{\beta}_{\boldsymbol{1}}$ is the weight of the pixel-wise luminance difference, $\boldsymbol{I}_{\boldsymbol{\Delta}}$, $\boldsymbol{\beta}_{\boldsymbol{2}}$ is the weight of phase-invariant structural similarity, $\boldsymbol{S}_{\boldsymbol{\Delta}}$, and $\boldsymbol{\beta}_{\boldsymbol{3}}$ is the weight of the interaction $\boldsymbol{I}_{\boldsymbol{\Delta}}\boldsymbol{S}_{\boldsymbol{\Delta}}$. To partially pool coefficient estimates across participants, the GLMM included participant and image combination as random effects.

| **Name** | **Estimate** | **SE** | **tStat** | **DF** | **pValue** |
| --- | --- | --- | --- | --- | --- |
| Intercept | 0.964 | 0.079 | 12.218 | 27657 | <.001 |
| $S_{\Delta}$ | -0.086 | 0.035 | -2.478 | 27657 | 0.013 |
| $I_{\Delta}$ | -0.035 | 0.035 | -1.016 | 27657 | 0.310 |

Table S.8: Formal model comparison. Here, we compare the three models described above, corresponding to the structural similarity-only model (S), the main effect model (M) including both structural similarity and pixel-wise luminance difference, and interaction model (I). Absolute ΔAIC, likelihood ratio statistics (LRStat), and *p* values are provided, calculated relative to the winning model (S).

| **Model** | **DF** | **ΔAIC** | **LRStat** | **pValue** |
| --- | --- | --- | --- | --- |
| S | 4 | 0 |  |  |
| M | 5 | 1.817 | 0.183 | .669 |
| I | 6 | 2.275 | 1.725 | .422 |
